# Mechanism of dual pharmacological correction and potentiation of human CFTR

**DOI:** 10.1101/2022.10.10.510913

**Authors:** Chi Wang, Zhengrong Yang, Blaine J. Loughlin, Haijin Xu, Guido Veit, Sergey Vorobiev, Oliver B. Clarke, Fan Jiang, Yaohui Li, Shikha Singh, Zachary Rich, Elizabeth R. Menten, Robert A. Grassucci, Wei Wang, Allison Mezzell, Ziao Fu, Kam-Ho Wong, Jing Wang, Diana R. Wetmore, R. Bryan Sutton, Christie G. Brouillette, Ina L. Urbatsch, John C. Kappes, Gergely L. Lukacs, Joachim Frank, John F. Hunt

## Abstract

Cystic fibrosis (CF) is caused by mutations in a chloride channel called the human Cystic Fibrosis Transmembrane Conductance Regulator (hCFTR). We used cryo-EM global conformational ensemble reconstruction to characterize the mechanism by which the breakthrough drug VX445 (Elexacaftor) simultaneously corrects both protein-folding and channel-gating defects caused by CF mutations. VX445 drives hCFTR molecules harboring the gating-defective G551D mutation towards the open-channel conformation by binding to a site in the first transmembrane domain. This binding interaction reverses the usual pathway of allosteric structural communication by which ATP binding activates channel conductance, which is blocked by the G551D mutation. Our ensemble reconstructions include a 3.4 Å non-native structure demonstrating that detachment of the first nucleotide-binding domain of hCFTR is directly coupled to local unfolding of the VX445 binding site. Reversal of this unfolding transition likely contributes to its corrector activity by cooperatively stabilizing NBD1 and the transmembrane domains of hCFTR during biogenesis.

**Summary:** Cryo-EM global conformational ensemble reconstruction has been used to characterize the mechanism-of-action of a breakthrough pharmaceutical that corrects fatal protein-folding and channel-gating defects in the human cystic fibrosis transmembrane conductance regulator (CFTR).

## INTRODUCTION

In 2019, the US FDA approved a breakthrough triple-drug combination therapy called Trikafta (Keating et al., 2018; Middleton et al., 2019) (VX445/VX661/VX770 or Elexacaftor/Tezacaftor/Ivacaftor) for the treatment of cystic fibrosis (CF), one of the most common fatal human genetic diseases. Clinical studies suggest that this treatment will extend the lifespans of ∼95% of CF patients by several decades. The development of Trikafta represents a milestone in molecular medicine that relied on synthesis of cutting-edge technologies in diverse areas of molecular biology and medicinal chemistry (Anderson et al., 1991; Cheng et al., 1990; Denning et al., 1992; Lukacs et al., 1994; Riordan et al., 1989). The identification in 1989 of the gene mutated in CF patients, the human Cystic Fibrosis Transmembrane Conductance Regulator (hCFTR), represented the first time that molecular biology methods were used to identify the cause of a genetic disease (Riordan et al., 1989). Cell biological and electrophysiological studies conducted during the ensuing decade demonstrated that hCFTR is an ATP-gated chloride channel (Anderson et al., 1991) that plays a central role in maintaining water homeostasis in the lungs and some secretory tissues. Cell biological studies furthermore established that the two most common CF mutations, F508del and G551D, cause different molecular defects. F508del (Cheng et al., 1990; Denning et al., 1992; Lukacs et al., 1994; Riordan et al., 1989) causes a biogenesis defect, reflecting destabilization of hCFTR, that leads to degradation of ∼99% of the protein by the quality-control apparatus in the endoplasmic reticulum (ER) prior to export to the site-of-function at the cell surface. G551D (Yang et al., 1993) instead prevents activation of the chloride channel in natively folded hCFTR that has reached the cell surface.

The insights generated by these fundamental studies led to the development of cell-based screens harnessing the green fluorescent protein (GFP) for empirical identification of small molecules enhancing mutant hCFTR biogenesis (folding correctors) (Van Goor et al., 2011; Van Goor et al., 2006) or channel activation (potentiators) (Gonzalez and Negulescu, 1998; Van Goor et al., 2009; Van Goor et al., 2006). This empirical approach led to the development of Trikafta (Keating et al., 2018; Middleton et al., 2019), but the mechanisms-of-action of the constituents of this breakthrough treatment remain unclear for VX661 (Fiedorczuk and Chen, 2022; Ren et al., 2013; Van Goor et al., 2011; Van Goor et al., 2006) and VX770 (Hadida et al., 2014; Liu et al., 2019; Van Goor et al., 2009) and completely unknown for VX445 (Veit et al., 2020; Veit et al., 2021)). We herein present structural and cell biological studies, supported by contemporaneously reported hydrogen-deuterium exchange mass spectrometry (HDX-MS) studies (Soya et al., Submitted), that identify the binding site of VX445 and provide insight into the molecular mechanisms underlying both its corrector and potentiator activities.

The hCFTR protein shares a common domain organization (Fig. 1A) and mechanochemical mechanism (Smith et al., 2002; Vergani et al., 2005) with proteins in the <u>A</u>TP-<u>B</u>inding <u>C</u>assette (ABC) Transporter superfamily (Allikmets et al., 1996; Chen, 2013; Mi et al., 2017). A pair of the stereotyped ATPase domains, from which the superfamily takes its name, function as an ATP-dependent mechanical clamp (Smith et al., 2002) that drives rearrangement of attached trans<u>m</u>embrane (TM) domains (Chen, 2013; Mi et al., 2017), which leads to formation of an internal chloride channel in the case of hCFTR (Aleksandrov et al., 2009; Aleksandrov et al., 2015; Vergani et al., 2005). Clamp closure is driven by encapsulation of two ATP molecules at the interface between two ABC domains (Smith et al., 2002), which are called <u>N</u>ucleotide-<u>B</u>inding <u>D</u>omains 1 and 2 (hNBD1/2) in hCFTR (Lewis et al., 2004; Protasevich et al., 2010; Yang et al., 2018). In this state, each ATP directly contacts the “Walker A” phosphate-binding motif (Walker et al., 1982) in one NBD and the ABC superfamily “Signature Sequence” in the other NBD (Smith et al., 2002). The G551D mutation (Veit et al., 2021; Yang et al., 1993) occurs in the Signature Sequence in hNBD1, which converts its canonical LSG<u>G</u>Q amino-acid sequence to LSG<u>D</u>Q. Gly551 in the wild-type (WT) sequence directly hydrogen-bonds (H-bonds) to the γ-phosphate of one ATP molecule in the closed state of the hNBD1/hNBD2 clamp in hCFTR (Zhang et al., 2017), so mutation to aspartate has been hypothesized to block channel opening by preventing closure of the clamp due to repulsive interaction with this ATP molecule (Bompadre et al., 2008).

**Figure 1.**
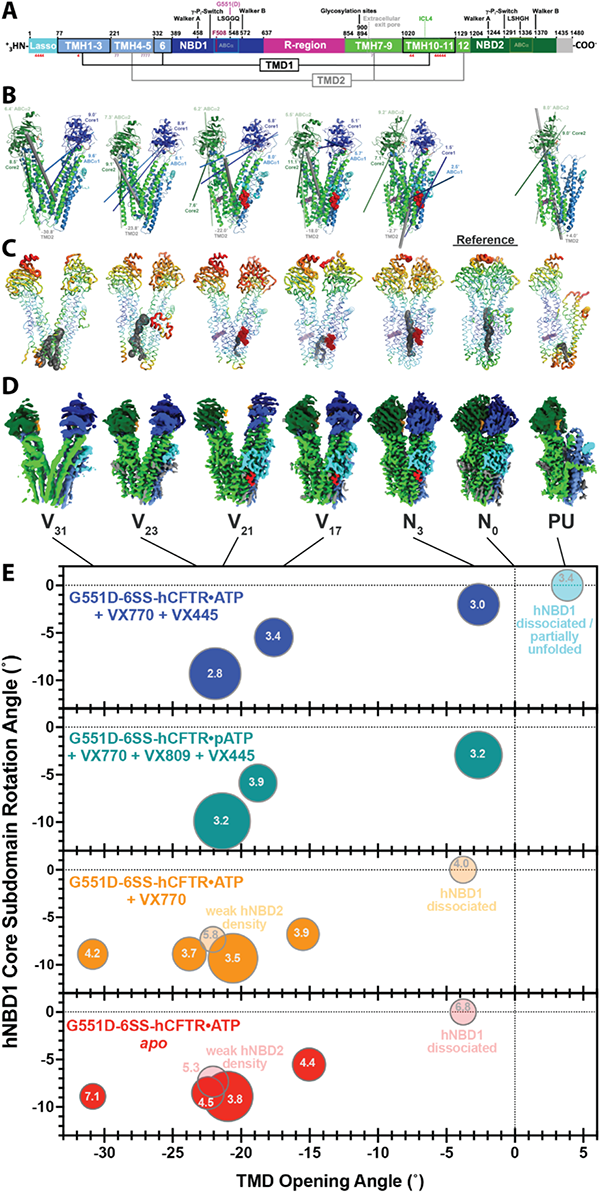
Structures and population distributions in global conformational ensemble reconstructions (GCERs) of G551D-6SS-hCFTR. (**A)** Linear sequence schematic illustrating domain organization, catalytic motifs, and drug-interaction sites. Each numeral under the bar represents one residue interacting with VX445 (“4” in red) or VX770 (“7” in dark violet). (**B)** Ribbon diagrams for the structures color-coded as in panel **A**, with VX445 and VX770 shown in stick representation in red or dark violet, respectively. The lines show the axes for the rotations of the subdomains compared to the PKA-phosphorylated E1371Q-6SS-hCFTR reference structure (labeled N_0_ in panels **D**-**E**), with the axes for the ABCa and Core subdomains in hNBD1 colored light and dark blue, respectively, the axis for TMD2 colored gray and shown with a wider diameter, and the axes for the ABCa and Core subdomains in hNBD1 colored medium and dark green, respectively. The structures are labeled according to the inter-TMD rotation angle relative to the reference structure, with V and N designating “<u>V</u>-shaped” *vs.* “<u>n</u>arrow” conformations, respectively. (The structures in the absence of VX445 with dissociated or weak NBDs are not shown in panels **A**-**D** even though their populations are reported in panel **E**.) (**C)** B-factor-encoded backbone representations of the reconstructed conformations using the default color ramp in PyMOL in which the most rigid backbone segments are thin and blue, while the most dynamic segments are thick and red. The dark gray spheres show the internal channels extending from the largest internal cavity located near the extracellular (bottom) surface, as calculated by CAVER (Pavelka et al., 2016). VX445 and VX770, when present, are shown in red or dark violet dot representation, respectively. The leftmost conformations both show substantial differences in the extracellular region of TMD2 (Fig. S6) that produce sizable packing voids that might be occupied by lipids. (**D)** EM density maps for each structure colored as in panels **A**-**B**. (**E)** GCER population distributions. The diameter of each circle represents the relative population of each reconstructed conformation, while the number inside the circle gives the gold-standard FSC resolution of that structure from non-uniform refinement in cryoSPARC (Punjani et al., 2020). The horizontal axis shows the rotation angle of the non-domain swapped helices in TMD2 (TMH7-9 & TMH12) relative to the N_0_ reference structure, while the vertical axis shows the rotation angle of the F1-like ATP-binding Core subdomain in hNBD1. The positive angle for the partially unfolded structure indicates a rotation narrowing rather than widening the inter-TMD1-TMD2 interface around an approximately vertical rotation axis with a nearly orthogonal orientation compared to the rotation axes in the V-shaped structures (as illustrated in **Movie 5**).

The hCFTR protein contains four additional structural modules – two TM domains (TMD1/2), a 67-residue N-terminal extension of TMD1 called the “Lasso” (Zhang et al., 2017), and an ∼200 residue regulatory (R) region (Ostedgaard et al., 2000; Riordan et al., 1989) (Fig. 1A). This last sequence module is predominantly disordered but local segments of it block channel opening by binding at the surfaces of TMD1/2 that combine to form the chloride channel upon closure of the ATP-dependent mechanical clamp (Zhang et al., 2017). Protein Kinase A (PKA) opens hCFTR channels by phosphorylating and thereby disordering these activity-inhibiting segments of the R region. Each TMD of hCFTR comprises a bundle of six TM α-helices (TMHs), but one pair of helices in each structural unit is domain-swapped, meaning formed by a discontinuous segment of primary sequence located between segments that form part of the other TMD. Therefore, TMD1 comprises TMHs 1, 2, 3, 10, 11, and 6, while TMD2 comprises TMHs 7, 8, 9, 4, 5, and 12 based on their order in the primary sequence (Fig. 1A). This domain-swapping has been proposed to be an obstacle to efficient biogenesis of hCFTR and to contribute to the defect caused by the F508del mutation (Fiedorczuk and Chen, 2022; Kleizen et al., 2021; Ren et al., 2013), which occurs at a site in hNBD1 that interacts directly with the protein segment connecting TMH10-11 in the domain-swapped region of TMD1. This segment and the cytosolic regions of TMH10-11 are called <u>I</u>ntra<u>c</u>ellular <u>L</u>oop 4 (ICL4).

## RESULTS

### Global Conformational Ensemble Reconstructions

To characterize the mechanism of potentiator action, we undertook cryogenic electron microscopy (cryo-EM) studies of G551D-hCFTR because this mutant protein requires potentiator binding for stable occupancy of the open-channel conformation. High-level production of G551D-hCFTR in mammalian tissue-culture cells (Fig. S1) was promoted by the introduction of a set of six previously characterized stabilizing mutations in hNBD1 (Yang et al., 2018). The PKA-phosphorylated, ATP-bound structure of this 6SS-hCFTR protein construct harboring the E1371Q mutation in the ATPase active site in hNBD2, which blocks ATP hydrolysis (Aleksandrov et al., 2015; Vergani et al., 2005), has no significant differences compared to the structure of WT hCFTR harboring the same hydrolysis-deficient mutation (Zhang et al., 2017) (Fig. S2). We performed cryo-EM reconstructions of PKA-phosphorylated G551D-6SS-hCFTR solubilized in digitonin and cholesterol-hemisuccinate (Zhang et al., 2017) (CHS) in the presence of ATP, ATP/VX770, ATP/VX770/VX445, or phenylethyl-ATP/VX770/VX809/VX445 (Fig. 1B-E). VX770 (Van Goor et al., 2014; Veit et al., 2021; Yu et al., 2012) and VX445 (Veit et al., 2021) both potentiate G551D-hCFTR channel activity, while VX445 additionally acts as a corrector that improves the biogenesis of hCFTR harboring F508del or a variety of other disease-causing mutations believed to destabilize the protein (Middleton et al., 2019; Veit et al., 2020). Phenylethyl-ATP (pATP) has been shown to have modest potentiator activity for G551D-hCFTR (Bompadre et al., 2008). VX770 was solubilized in digitonin for delivery to hCFTR samples at a defined concentration (∼5 µM free VX770). However, adding VX445 directly to purified hCFTR samples yielded problematic background in cryo-EM imaging, so we developed a procedure (explained in Fig. S3) using defatted bovine serum albumin to deliver VX445 to hCFTR at a roughly stoichiometric level.

We attempted to establish the approximate population of all hCFTR conformations present in our cryo-EM samples. We have dubbed this emerging approach (Lutomski et al., 2022; Wang and Boudker, 2020) in single-particle reconstruction studies <u>G</u>lobal <u>C</u>onformational <u>E</u>nsemble <u>R</u>econstruction (GCER). It builds on the “story-in-a-sample” approach (Agirrezabala et al., 2012; Fischer et al., 2010; Frank, 2013) enabled by the development of direct electron detectors by harnessing improved classification and reconstruction algorithms developed during the past decade (Kimanius et al., 2021; Punjani and Fleet, 2021; Punjani et al., 2020; Scheres, 2012) to produce more qualitatively complete and quantitatively reliable characterization of the conformational ensemble in a protein sample.

Following promiscuous template-picking with projections from the widest and narrowest hCFTR conformations filtered to 20 Å, only limited cleaning was performed of 2-dimensional classes, retaining all classes roughly matching the dimensions of hCFTR to avoid conformational bias. Following procedures to try to identify all distinct protein conformations present in each sample (see Methods), particles were simultaneously cleaned and assigned to conformational classes by iterative cycles of heterorefinement in cryoSPARC (Punjani et al., 2020) against reference volumes representing every distinct hCFTR conformation we have reconstructed together with six dummy volumes ranging in size from an empty digitonin micelle to that of hCFTR (Li et al., 2022). Particles assigned to hCFTR volumes were passed to the subsequent cycle, while those assigned to dummy volumes were discarded. After at least 3 cycles, single-and multi-class *ab initio* reconstruction and 3-dimensional variability analyses (Punjani and Fleet, 2021) (3D-VA) were performed on the particles assigned to each hCFTR volume to determine whether additional conformations could be detected. Volumes from particles that did not yield an *ab initio* reconstruction with an equivalent structure and resolution to heterorefinement were discarded, and only one was retained when multiple volumes had qualitatively equivalent structures. All of the resulting structurally distinct classes were used as references to initiate at least six cycles of iterative heterorefinement starting again with the original particle stack obtained from limited 2-dimensional cleaning (Li et al., 2022). The relative populations assigned to each class are stable during this procedure and similar in all particles or those with >99% probability of belonging to each class after the final cycle (Fig. S4). The resolutions of the individual structures in the GCERs range from 2.8-7.1 Å with most being better than 3.8 Å resolution (Fig. 1E and Table S1).

### G551D-hCFTR dynamically interconverts between diverse conformational states

All hCFTR samples we have imaged in digitonin/CHS yield reconstructions of multiple qualitatively distinct protein conformations at near-atomic-resolution (Fig. 1 and data not shown), including not only G551D-6SS-hCFTR samples but also samples of constructs yielding nearly 100% open probability in membranes (*e.g.*, E1371Q-6SS-hCFTR (Aleksandrov et al., 2015)). The conformational diversity in the GCERs from those constructs indicate that digitonin/CHS biases hCFTR away from the open-channel conformation compared to native phospholipid membranes. We have categorized the observed G551D-6SS-hCFTR conformations based on their inter-TMD rotation angles (*i.e.*, <u>V</u>-shaped conformations V_31_, V_21_, V_23_, and V_17_ without an internal chloride channel and <u>n</u>arrow conformation N_3_ with an internal chloride channel) compared to the “N_0_” ATP-bound E1371Q-6SS-hCFTR reference conformation with an internal chloride channel (Fig. 1B-E).

While a sizable majority of the population in every sample comprises particles in which TMD1/hNBD1/TMD2/hNBD2 are well-ordered, all but the pATP/VX770/VX809/VX445 sample unambiguously contain a significant population of molecules (9-15%) with minimal density for hNBD1, which is either diffusing in a detached state or potentially unfolded (Protasevich et al., 2010). As discussed below, these hNBD1-dissociated structures show at least partial unfolding of domain-swapped TMH10-11/ICL4. Some samples also contain a smaller population with very weak hNBD2 density and a shifted conformation in the interacting loops in TMD2 (Fig. S5), indicating the domain is either highly dynamic or partially dissociated. The conformations with TMD1/hNBD1/TMD2/hNBD2 well-ordered likely represent a continuum of dynamically interconverting conformational states prior to vitrification for cryo-EM imaging, and they also likely interconvert with the NBD-dissociated conformations, although, as discussed below, some hNBD1-dissociated molecules seem likely to be kinetically trapped in a partially unfolded state.

Performing 3D-VA after combining particles from different hNBD1-containing structures provides a qualitative illustration of the overall conformational dynamics of hCFTR in digitonin/CHS (**Movie 1**). This analysis shows two distinct large-scale tweezer-like motions of the TMDs, one along the conformational trajectory that mediates channel formation. These TMD motions are coupled to several modes of NBD motion involving differential rotation of the α-helical (ABCα) subdomain (Karpowich et al., 2001), which contains the ABC Signature Sequence, relative to the F_1_-ATPase-like ATP-binding Core (Karpowich et al., 2001), which contains the Walker A motif (Walker et al., 1982) (Fig. 1A-B). The ABCα subdomain and ATP-binding Core in each NBD are connected by a 7-residue linker called the γ-phosphate switch (Karpowich et al., 2001) or Q-loop (Fig. 1A) because it contains a conserved glutamine that contacts the γ-phosphate of ATP in the closed, pre-hydrolysis state of the ATP-dependent mechanical clamp formed by the NBDs. The ABCα subdomain and ATP-binding Core have been demonstrated to maintain rigid internal conformations but rotate relative to one another in structures of isolated hNBD1 from hCFTR (Lewis et al., 2010) as well as in different functional conformational states of homologous domains from ABC superfamily proteins (Karpowich et al., 2001). Equivalent rigid-body inter-subdomain rotations, which mediate ATP-dependent allosteric conformational changes in many of those proteins (Chen, 2013), contribute to the observed conformational dynamics of hCFTR and also activation of the G551D mutant byVX445 (Fig. 1B and **Movies 1-3**).

Performing 3D-VA on the particles used to reconstruct any individual G551D-6SS-hCFTR structure shows qualitatively similar dynamics (**Movies 2-3**) but on a smaller scale than in the combined particle sets including multiple structures in the GCER (**Movie 1**). Therefore, the individual structures represent clusters of molecules in similar but not identical conformations within the global conformational ensemble. The 3D-VA movies from the individual structures still show some degree of tweezering between TMD1/TMD2 as well as more prominent rotations of the ABCα subdomains and ATP-binding Cores in both NBDs. Refined atomic displacement parameters (B-factors) for these structures reflect these qualitative differences in dynamics in different regions (Fig. 1C). Notably, B-factor analysis shows minimal conformational variation in our E1371Q-6SS-hCFTR/ATP N_0_ reference structure (Fig. 1C). This structure is nearly indistinguishable from the previously reported E1371Q-hCFTR/ATP structure (Zhang et al., 2017) (Fig. S2), both of which have the mechanical clamp formed by hNBD1/hNBD2 in the canonical ATP-bound, closed conformation of the homologous domains observed in many ABC superfamily proteins (Chen, 2013; Smith et al., 2002). Therefore, the G551D mutation prevents adoption of the dynamically stable state that represents the predominant conformation found in hCFTR constructs without G551D under equivalent biochemical conditions, which has been interpreted to represent the open, high-chloride-conductance conformation of hCFTR (Zhang et al., 2017).

All structures exhibiting density for both NBDs in the PKA-phosphorylated G551D-6SS-hCFTR/ATP ensemble in the absence of potentiator (V_17_, V_21_, V_23_, and V_31_ — bottom of Fig. 1E) have widely separated NBDs and TMDs compared to the N_0_ reference structure without the G551D mutation. The 17°-31° inter-TMD rotation physically disrupts the chloride-conductance channel formed at their interface (Fig. 1C), likely stopping ion conductance. Compared to the V_17_ and V_21_ conformations, the V_23_ and V_31_ conformations both show refolding of the C-terminus of TMH5 coupled to unfolding of the N-terminus of TMH8 (Fig. S6), which disrupts the VX770 binding site (see below) and creates packing voids in TMD2 (Fig. 1C) that may be filled with lipids. The cryo-EM maps clearly show unhydrolyzed ATP in the ATP-binding sites in both NBDs in these structures (*e.g.*, as shown in Fig. 5 below). Therefore, our GCER ensemble strongly supports the hypothesis (Bompadre et al., 2008) cited above that the disease-causing G551D mutation blocks hCFTR channel activation by preventing the mechanical clamp formed by its NBDs from adopting the canonical ATP-bound, closed conformation observed in the hCFTR N_0_ reference structure and other ABC superfamily proteins.

### Visualization of a lipid raft that may regulate hCFTR channel gating

Our GCER for G551D-6SS-hCFTR/ATP includes one V_21_ structure containing ∼20 well-ordered cholesterol or phospholipid molecules organized in a roughly hexagonal packing geometry characteristic of a lipid raft (Levental et al., 2020) (Fig. 2). This putative raft, which is bound to the surface of TMD2 at a site located in the outer-leaflet of the plasma membrane, is present in only one of seven structural classes reconstructed for G551D-6SS-hCFTR/ATP. The GCER contains a second V_21_ structure with an equivalent protein conformation but density for only ∼5 cholesterol molecules in the raft that directly contact the surface of TMD2 (Fig. 2). The two V_21_ conformational classes have roughly equal size and together and account for 36% of the population (Figs. 1E and **S4**), more than twice that of any other class. Notably, the surface-bound cholesterol molecules retained in the V_21_ structure without the raft are not observed in hCFTR structures in which the internal chloride channel is formed (*e.g.*, the N_3_ conformation shown on the right in Fig. 2), suggesting that lipid raft binding at this site could potentially inhibit/regulate channel activation.

**Figure. 2:**
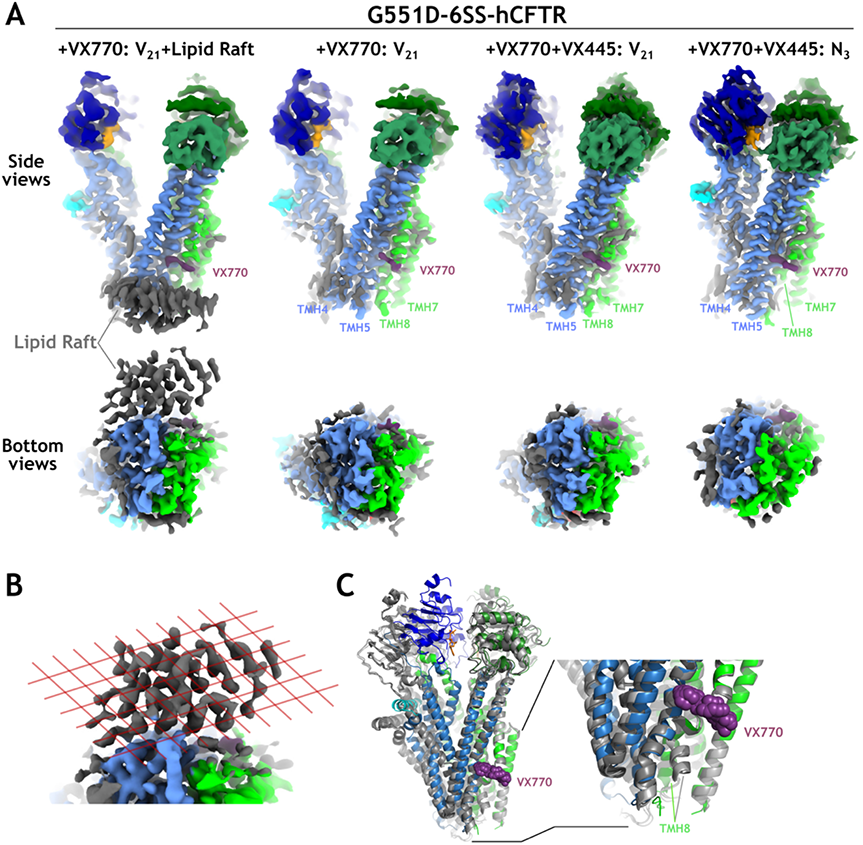
The N-terminal segment of TMH8 contributing to the exit pore of the hCFTR chloride channel changes conformation in the N_3_ vs. V_21_ conformations but not upon lipid raft binding to the V_21_ conformation. **(A)** EM density maps with protein domains colored as in Fig. 1A and lipid species colored gray. The density suggests these species are primarily but not exclusively cholesterol, with some likely being diacylphosophilids or potentially fatty acid segments. **(B)** Zoomed-in view of the putative lipid raft on the left in panel **A** with semi-transparent red lines illustrating a 2-dimensional hexagonal lattice (P6 plane group symmetry). **(C)** Ribbon diagrams showing superposition of the refined models for all the maps shown in panel **A** based on least-squares alignment of the non-domain-swapped helices in TMD2. TheV_21_ structures, which have nearly indistinguishable conformations, are shown in different shades of gray, while the N_3_ structure is colored by domain organization as in Fig. 1A. The N_3_ structure shows a conformational shift in two segments contacting the lipid raft observed in the outer leaflet of the bilayer in the V_21_ structure (*i.e.*, in the N-terminus of TMH8 and the C-terminus of TMH5 directly below the VX770 binding site in the orientation depicted here).

### VX445 but not VX770 strongly modulates hCFTR conformation in digitonin/CHS

Addition of VX770 alone to G551D-6SS-hCFTR/ATP produces minimal changes in the conformational distribution determined by GCER (second graph from the bottom in Fig. 1E) even though three of the six distinct protein conformations (V_17_, V_21_, and hNBD1-dissociated) show clear EM density for the drug at the previously identified VX770 binding site (Liu et al., 2019) near the center of the bilayer in TMD2 (Fig. 3A and data not shown). Two of the remaining conformations (V_23_ and V_31_) show no density for the drug, presumably because they exhibit substantial conformational changes at the VX770 binding site involving displacement of the N-terminus of TMH8 by the C-terminus of TMH5 (Fig. S6), while the final conformation with weak hNBD2 density is too low in resolution to reliably assess whether the drug is bound (Fig. S5). Therefore, despite convincing observation of VX770 binding in three of five conformations reconstructed at near-atomic resolution (Fig. 1E), the drug does not produce a detectable change in the conformational distribution of G551D-6SS-hCFTR in digitonin/CHS. Additional GCERs that we have performed on 6SS-hCFTR and F508del-6SS-hCFTR in digitonin/CHS similarly show no detectable change in the conformational ensembles upon binding of VX770 (manuscript in preparation). The failure to observe a significant conformational change upon binding of VX770, which efficiently opens G551D and F508del channels in membranes (Yeh et al., 2017), suggests phoshoplipid interactions with hCFTR play a critical role in the mechanism-of-action of this potentiator.

**Figure. 3:**
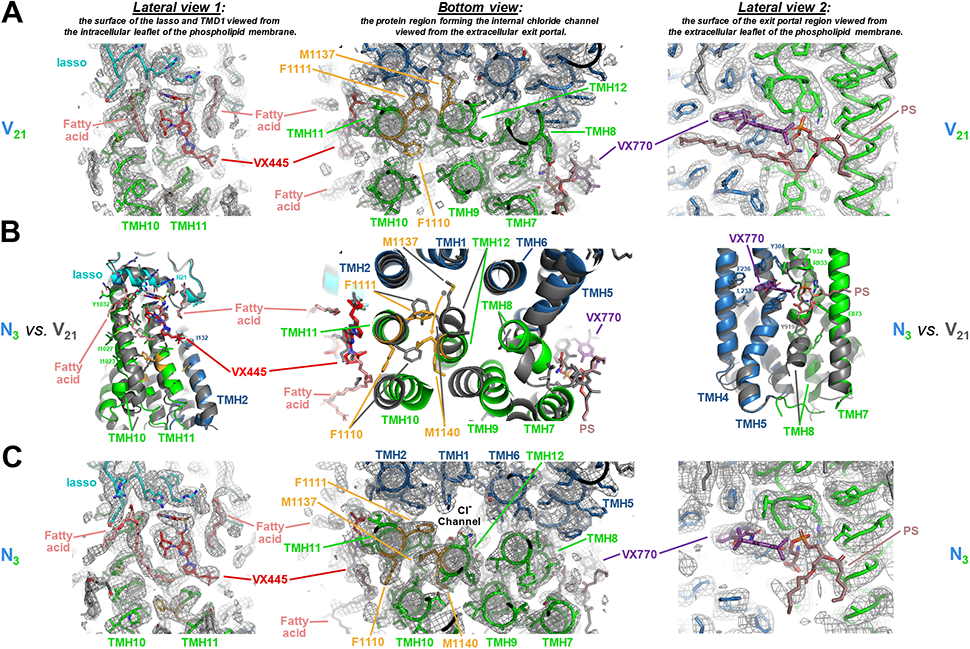
Structural changes in the VX445 binding-site in the V_21_ vs. N_3_ conformations of hCFTR. **(A,C)** Three views of cryo-EM density maps (gray mesh) for the G551D-6SS-hCFTR/ATP/VX770/VX445 V_21_ (panel **A**) and N_3_ (panel **C**) structures superimposed on coordinate models colored according to domain organization as in Fig. 1A. **(B)** Ribbon diagrams of the coordinate models for the V_21_ (backbone and carbon atoms in gray) and N_3_ structures (backbone and carbon atoms colored according to domain organization except for residues F1110, F1111, M1137, and M1140, which are colored orange to highlight their positions). The carbon atoms in VX770 and the associated putative inverted phosphatidylserine (PS) are colored in purple and violet, respectively while the carbon atoms in VX445 and associated phospholipids are colored in red and pink, respectively. Oxygen atoms are colored red, nitrogen atoms are colored blue, and sulfur atoms are colored yellow in all molecules. The protein structures were aligned in the non-domain-swapped core of TMD1.

An unidentified EM density feature has previously been observed immediately adjacent to the VX770 binding site (Liu et al., 2019), which occurs at a mid-membrane break in TMH8 that leads to exposure of peptide bonds from residues L926-F932 to the phospholipid bilayer (Fig. 3 right). This highly unusual structural feature creates a polar cavity in hCFTR in the hydrophobic core of the bilayer directly adjacent to the binding site for the strongly apolar compound VX770. This polar cavity is directly above the N-terminus of TMH8, which forms part of the gate of the hCFTR chloride channel in the extracellular leaflet of the membrane. Our best reconstructions of the VX770 binding site suggest the unidentified density feature may be the headgroup of a phosphatidylserine (PS) molecule adopting an inverted configuration in the outer leaflet of the membrane (Fig. 3 right). PS has been shown to greatly increase the thermal stability of purified hCFTR (Hildebrandt et al., 2017). The negatively charged phosphate group of the putative inverted PS molecule is located immediately beneath the positively charged end of the helix dipole at the N-terminus of the second segment of TMH8 following its mid-membrane break, where it putatively hydrogen-bonds (H-bonds) to the hydroxyl group of VX770 and the sidechain of R933 as well as the protein backbone at the break site. Its serine group putatively H-bonds to the sidechains of residues E873 and Y919 as well the protein backbone at the break site (Fig. 3B right). The failure of VX770 to open hCFTR channels in our cryo-EM samples could reflect disruption by digitonin of energetically significant interactions between VX770 or this putative PS molecule with other phospholipids and the nearby gating region at the N-terminus of TMH8 in the extracellular leaflet of the membrane.

In contrast, addition of VX445 (Keating et al., 2018; Middleton et al., 2019; Veit et al., 2020; Veit et al., 2021) to G551D-6SS-hCFTR/ATP/VX770 produces a large change in conformational distribution (Fig. 1E) including appearance of a new conformation (N_3_) in which TMD1/TMD2 move inward by ∼14-18° to form most of the internal chloride channel observed in the N_0_ reference structure without G551D (Fig. 1C). The conformation of its cytoplasmic entrance is somewhat different in the N_3_ conformation than in the ATP-bound N_0_ reference structure that has been interpreted to represent the high-conductance state of hCFTR (Zhang et al., 2017), but its extracellular exit pore and most of its internal vestibule have equivalent conformations (Fig. 1C). Therefore, its electrophysiological properties could be similar given that the conductance-limiting region of the channel is believed to be the extracellular exit pore (Linsdell, 2017). Induction of a conformation with an internal chloride channel *in vitro* is consistent with the observation that VX445 acts as independent potentiator of G551D-hCFTR chloride-channel conductance in cells as well as a co-potentiator in combination with VX770 (Veit et al., 2021).

The V_21_, V_17_, and N_3_ structures in the G551D-6SS-hCFTR/ATP/VX770/VX445 GCER (Fig. 1B-E) all show VX445 bound in a membrane-embedded cavity in TMD1 (Figs. 3 left and center and **4A-B**) lined by the N-terminus of the Lasso (S18, R21, L24, R25) and segments of TMH2 (I132), TMH10 (L1029, Y1032), and TMH11 (W1098, R1102, M1105, I1106, V1108, I1109). However, there is no evidence in this GCER of the three widest conformations observed in the reconstructions in the absence of VX445 (V_31_, V_23_, and weak hNBD2 density in Fig. 1E). These structures all show weak or no density for the N-terminus of the Lasso (Figs. S5-S6), likely preventing VX445 binding given the contribution of this protein segment to the observed binding site, and this effect is likely to explain the reduction in their population in the presence of the drug given that its high binding energy (∼-10 kcal/mol) will stabilize binding-competent conformations compared to non-interacting conformations. The failure to observe the widest conformations in the presence of VX445 reinforces the conclusion that its binding shifts the conformational ensemble towards states with closer association of TMD1/TMD2, including the N_3_ state with the internal chloride channel formed at their interface (Fig. 1C-E). Therefore, VX445 binding to G551D-6SS-hCFTR solubilized in digitonin/CHS produces a clear shift in the population towards the open-channel conformation, while VX770 binding does not.

### Structural basis of hCFTR channel potentiation by VX445

Examining a superposition of the V_21_ and N_3_ structures with bound VX445 (Fig. 3B) shows the stereochemistry of its binding cavity changes in the N_3_ conformation in which the internal chloride channel is formed (Fig. 3C), while it is equivalent in the V_21_ (Fig. 3A) and V_17_ (data not shown) conformations without an internal chloride channel. The observed conformational differences can explain the drug’s potentiator activity assuming the free energy of the VX445-N_3_ complex is more favorable than the VX445-V_21_/VX445-V_17_ complexes. The conformational differences include Ångstrom-scale backbone shifts in different directions throughout the binding site and also in the position of VX445 relative to residues W1098-I1109 in TMH11 that form the core of its observed binding site (Fig. 3B left). The largest structural change occurs in TMH10, which shifts outward (as oriented in the left panels in Fig. 3) by ∼1.4 Å at Y1032 but ∼2.9 Å at I1023. These displacements propagate in a pendulum-line manner towards the N-terminus of TMH10, which moves by ∼7 Å, directly displacing the C-termini of TMH7/TMH9 and the N-terminus of TMH12, and these changes are accompanied by a substantial repacking of the interface between TMH12 and residues F1110-F1111 immediately adjacent to the VX445-interacting segment in TMH11 (Fig. 3B left and center). The region of TMH12 involved in this repacking directly contacts the N-terminus of TMH8, which forms part of the extracellular exit portal of the chloride channel, and this region of TMH8 also contacts the segments of TMH7/TMH9 shifted by the pendulum-like motion of TMH10 (Fig. 3B center), providing multiple pathways of allosteric structural interaction between the VX445 binding site and a protein segment likely to control chloride conductance.

The N_3_ conformation shows strong EM density for a fatty acid tightly sandwiched between VX445 and residues A1025-L1029 in TMH10, while the density for this fatty acid is substantially reduced in the V_21_ conformation (left in Fig. 3A *vs.* Fig. 3C), suggesting it contributes to the differential stabilization of the N_3_ conformation by the drug. Therefore, lipid interactions seem likely to make critical contributions to the mechanisms-of-action of both VX445 and VX770.

The NBDs in the narrowest N_3_ conformation of G551D-6SS-hCFTR/ATP/VX770/VX445 are clearly not in the same ATP-bound conformation observed in the N_0_ reference structure (Fig. 5), which matches the canonical closed state of the ATP-dependent mechanical clamp observed for the homologous domains in many ABC superfamily structures (Chen, 2013; Smith et al., 2002). Both of the ATP-binding sites are pulled apart in the narrowest N_3_ conformation of G551D-6SS-hCFTR/ATP/VX770/VX445 (Fig. 5A), although, surprisingly, substantially wider at the composite site formed by the Walker A in the core of hNBD1 and the Signature Sequence in the ABCα subdomain of hNBD2 (16.7 *vs.* 11.4 Å) than at the composite site formed by the Walker A in the core of hNBD2 and the mutated G551D-containing Signature Sequence in the ABCα subdomain of hNBD1 (12.1 *vs.* 10.1 Å). The unexpected observation of closer apposition of the NBDs at the site harboring the G551D mutation, even though it will electrostatically repel the ATP molecule at their interface, indicates that NBD proximity at this site is disproportionately important in allosterically stabilizing the TMD conformation with an internal chloride channel.

The N_3_ conformation of G551D-6SS-hCFTR/ATP/VX770/VX445 shows diffuse EM density for the mutated LSGDQ Signature Sequence harboring the G551D mutation in the ABCa subdomain of hNBD1 compared to both adjacent regions in that structure and the equivalent segment in the N_0_ reference structure (Fig. 5B), indicating elevated local dynamics at the mutation site. In the canonical ABC superfamily conformation represented by the N_0_ reference structure, the Signature Sequence forms an α-helical capping structure that encapsulates the γ-phosphate of ATP with its backbone amide groups and serine sidechain, while the glutamine at its C-terminus H-bonds to the ribose of ATP (right in Fig. 5) (Smith et al., 2002). The dominant conformation observed in our G551D-6SS-hCFTR/ATP/VX770/VX445 N_3_ structure instead shows the Signature Sequence harboring the G551D mutation shifted 4-5 Å away from the γ-phosphate of ATP, disrupting all of the specific H-bonds characteristic of the canonical structure, and the residues adjacent to its N-terminus making a limited number of non-canonical contacts to the ribose and adenine of the ATP. In the narrowest conformation observed in our GCER for G551D-6SS-hCFTR/pATP/VX770/VX809/VX445, which contained a 60-fold ratio of pATP:ATP, the fiducial sites in the Walker A in hNBD2 and the mutated LSGDQ Signature Sequence in hNBD1 have a similar separation as in the N_0_ reference structure (right in Fig. 5A), and the residues adjacent to the N-terminus of the Signature Sequence maintain somewhat more canonical contacts to the ribose and base of the pATP, which shifts slightly due to the phenylethyl-derivatization of its base. However, an ∼6.5° rotation of the ABCα subdomain containing the G551D mutation in hNBD1 again disrupts all interactions between its mutated LSGDQ Signature Sequence and the γ-phosphate of ATP (Fig. 5A). The diffuse EM density and non-canonical interactions in both the ATP and pATP structures (Fig. 5) indicate a lack of energetically stabilizing interactions between the NBDs at the composite ATP-binding site harboring the G551D mutation, while they are too far apart to interact at all at the other composite ATP-binding site.

We conclude that VX445 binding drives the TMDs into a conformation that pulls the NBDs together at the composite ATP-binding site harboring the G551D mutation but that the mutation prevents formation of stable stereochemical interactions across the hNBD1/hNBD2 interface at this site. Therefore, in contrast to the mechanochemical activation mechanism of WT hCFTR in which high-affinity ATP binding at this site pulls the NBDs together and they pull the TMDs into their active conformation, VX445 binding effectively reverses this pathway of allosteric communication by pulling the TMDs together and thereby driving the NBDs together despite their inability to interact stably at the G551D composite ATP-binding site due to the mutation. Therefore, VX445 binding induces allosteric communication along the same structural pathway but in the opposite direction, a conclusion strongly supported by recent HDX-MS studies (Soya et al., Submitted).

### Mutagenesis supports the observed VX445 binding site mediating correction and potentiation

Biogenesis experiments in immortalized human CF Bronchial Epithelial (CFBE) cells support VX445 binding to the site identified in our cryo-EM structures being responsible for its corrector activity (Fig. 4C-D). These experiments employed the disease-causing L206W mutation (Veit et al., 2020), which is located at a site in TMD1 ∼30 Å away from the VX445 binding site. L206W is correctable by VX445, VX661, and VX809 (Lumacaftor). The latter two correctors bind to the same site in TMD1 at the C-terminus of the Lasso ∼30 Å away from the VX445 binding site and ∼20 Å away from L206 (Fiedorczuk and Chen, 2022), and their activities are redundant with each other but additive with VX445 (Veit et al., 2020). We compared the sensitivity of hCFTR variants with mutations in the observed VX445 binding site to correction by VX809 *vs.* VX445 to compensate for non-specific structural perturbations (Fig. 4C-D). The R21G, R25S, and I1109F mutations attenuate correction by VX445 substantially more than by VX809, while M1105F specifically attenuates correction by VX445 without affecting correction by VX809. These results support the observed VX445 binding site being responsible for its corrector activity, a conclusion further supported by recently reported HDX-MS studies (Soya et al., Submitted).

**Figure 4.**
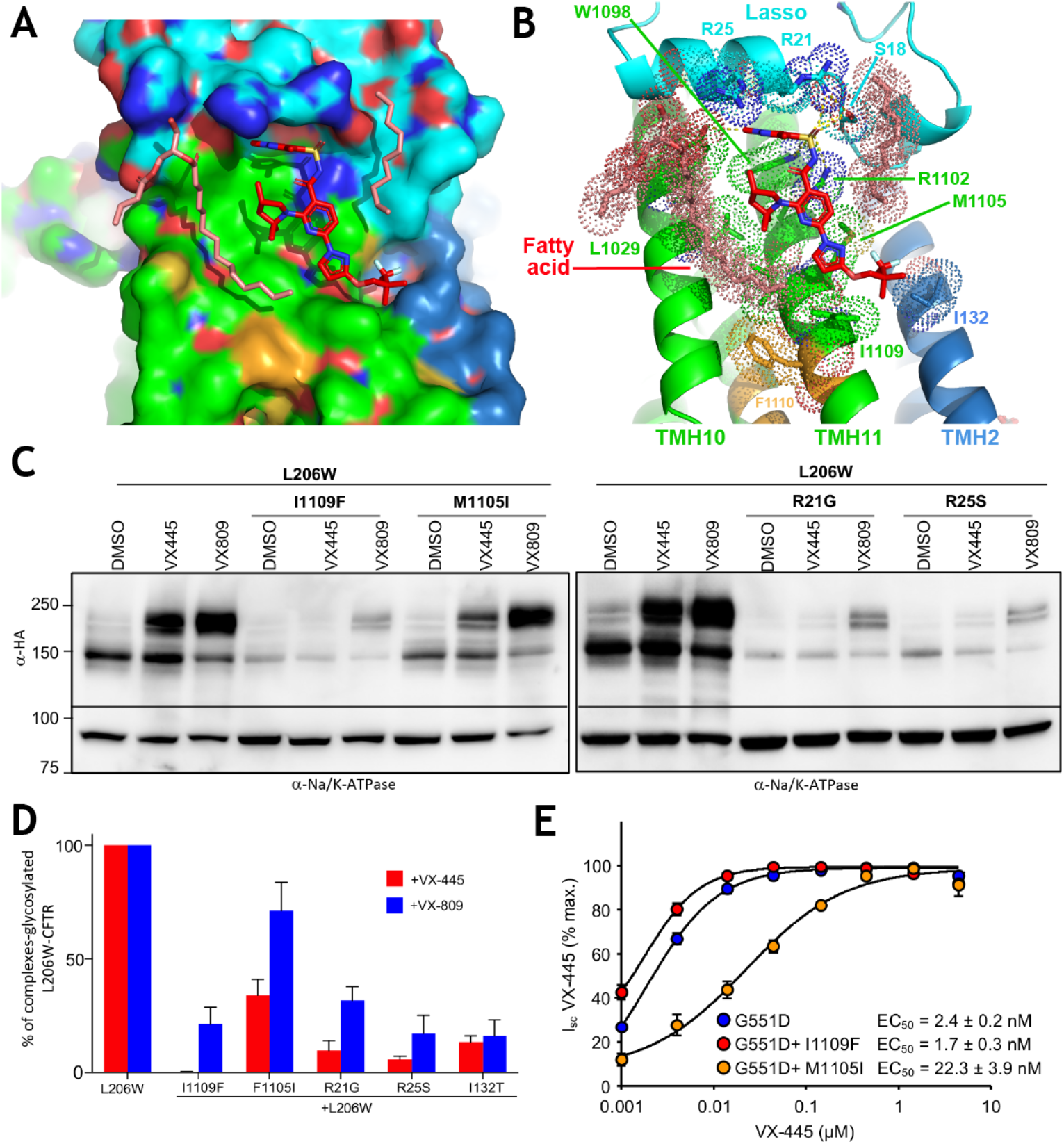
Structure and mutational analysis of the VX445 binding site in hCFTR. **(A-B)** Surface representation (panel **A**) and ribbon diagram (panel **B**) of the VX445 binding site in the N_3_ conformation of G551D-6SS-hCFTR/ATP/VX770/VX445 with the protein backbone and carbon atoms colored according to domain organization as in Fig. 1A, except for the carbon atoms in residues F1110, F1111, M1137, and M1140, which are colored orange. The carbon atoms in VX445 and adjacent phospholipid fragments are colored red and pink, respectively. The sidechains and lipid molecules contacting VX445 are shown with dot surfaces. **(C)** Immunoblots showing effect of mutations in the VX445 binding site on expression and maturation of L206W-CFTR-3HA in CFBE41o-immortalized human bronchial epithelial cells grown overnight in the absence or presence of VX-809 (3 mM) or VX-445 (2 mM). Equal amounts of lysate were loaded on the gels. **(D)** Quantification of the results in panel **C** using the lower immunoblots with an antibody against Na^+^/K^+^-ATPase to normalize for cell mass. **(E)** Effect of mutations in the VX445 binding site on the VX445 co-potentiator effect on forskolin-stimulated and VX770-stimulated short-circuit currents in hCFTR-expressing CFBE41o-cells. Error bars in panels **G**-**I** represent the standard error of the mean for 3 biological replicates.

Electrophysiology experiments in the same cells show that the I1109F mutation significantly increases the concentration of VX445 needed to potentiate G551D-hCFTR conductance, while the M1105I mutation does not (Fig. 4E), although it does attenuate the VX445-stimulated conductance level compared to the VX770-stimulated level (Fig. S7). Differential influence of mutations on corrector *vs.* potentiator activities is consistent with the observed conformational shift in the VX445 binding site (Fig. 3 left and center) in the activated N_3_ conformation compared to inactive V_21_ conformation, which is likely to be dominant during biogenesis in the ER, because the same stereochemical perturbation is likely to have different energetic effects when the structure of the drug interaction site changes. For example, the fatty acid cofactor that interacts with VX445 in the N_3_ conformation (Figs. 3B-C left and 4A-B) is shifted and shows substantially lower density in the V_21_ conformation (Fig. 3A-B left), suggesting it is likely to be important for potentiation but not correction. Therefore, the electrophysiology data in Figs. 4E and S7 support the VX445 binding site observed in our cryo-EM structures also being responsible for its potentiator activity.

### 3.4 Å reconstruction of an hCFTR conformation with the VX445 binding site unfolded

The GCER for G551D-6SS-hCFTR/ATP/VX770/VX445 includes a 3.4 Å hNBD1-dissociated and <u>p</u>artially <u>u</u>nfolded (PU) structure (Fig. 6 and **Movies 4-5**) that provides insight into the molecular mechanism by which VX445 binding corrects the biogenesis defect caused by numerous disease-causing mutations including F508del (Fig. 7). This structure (Fig. 6C-D) lacks EM density for the N-terminus of the Lasso, the entirety of TMH10, the cytoplasmic region of TMH11, and ICL4, which connects TMH10-11 and mediates most packing contacts between TMD1 and hNBD1, including all direct contacts to F508. Notably, recently reported HDX-MS results show that mutations that impair hCFTR biogenesis dynamically destabilize these protein segments more than others and that corrector drugs specifically offset these effects (Soya et al., Submitted). The PU cryo-EM map shows diffuse density in the TM region of TMH11 and even more diffuse density extending from the N-terminus of that segment roughly along the surface of the digitonin micelle, which is likely to represent a weakly-ordered alternative conformation of the cytoplasmic segment of TMH11. There is a similarly diffuse extension embedded in the surface of the digitonin micelle near the C-terminus of TMH9 that is likely to represent weakly-ordered alternative conformations of the N-terminal segment of TMH10 (Fig. 6D and **Movie 4**). The ability to reconstruct this partially unfolded hCFTR structure at near-atomic resolution indicates that binding of neither the domain-swapped TMH10-11 nor hNBD1 is required for stable folding of the core of TMD1.

Most residues forming the VX445 binding site are disordered in this PU conformation (Fig. 6D and **Movies 4-5**). Moreover, hNBD1-dissociated structures we have determined in GCERs from 11 hCFTR samples containing different protein constructs or nucleotides/drugs all show reduced or no density for the N-terminus of the Lasso and TMH10-11 (Figs. 1B-D and **6B-D** and data not shown). These observations support unfolding of these segments, which form most of the VX445 and hNBD1 binding sites on TMD1, being strongly thermodynamically coupled to hNBD1 dissociation. Therefore, VX445 binding will simultaneously stabilize the folded conformation of these segments and the bound state of hNBD1. We hypothesize this thermodynamic effect explains the corrector activity of VX445 during hCFTR biogenesis (Fig. 7 center).

Persistence of the PU conformation in the presence of VX445 in our G551D-6SS-hCFTR/ATP/VX770/VX445 sample (upper right in the top panel of Fig. 1E) suggests that this hNBD1-dissociated structure (Fig. 6C-D and **Movies 4-5**) is kinetically trapped in the unfolded state, at least in digitonin micelles. Notably, the protein sample used to produce this GCER contained a higher level of immaturely glycosylated “Band B” relative to the properly matured “Band C” compared to most of our hCFTR protein preparations (Fig. S1C *vs.* S1A), suggesting the possibility the PU structure could potentially represent an ER-resident biogenesis intermediate.

Our GCERs on 12 hCFTR samples (Fig. 1 and data not shown) suggest that the level of the kinetically-trapped, hNBD1-dissociated PU species varies in different preparations and that most samples contain a mixture of this species and a conformation with hNBD1 reversibly dissociated and the intracellular segments of TMH10-11, including ICL4, reversibly unfolded. The NBD1-dissociated structures from our samples without VX445 (Fig. 6B) show significant differences from the kinetically-trapped PU structure (Fig. 6C-D) consistent with the former containing molecules with hNBD1 reversibly dissociated. These maps show more residual density for the native conformation of TMH10-11, especially in the membrane-embedded region of TMH10, and a somewhat different packing angle between TMD1/2. Given the difficulty in classifying particles the size of hCFTR with relatively small conformational differences, these maps could represent a mixture of hCFTR molecules with hNBD1 reversibly dissociated and in the kinetically-trapped PU state. The high-quality of the reconstruction of the kinetically-trapped state in the G551D-6SS-hCFTR/ATP/VX770/VX445 sample could reflect the corrector activity of VX445 reducing the population of molecules with hNBD1 reversibly dissociated (Fig. 7), yielding a cleaner reconstruction of the kinetically-trapped PU state because a higher proportion of hNBD1-dissociated molecules are in that conformation rather than the reversibly dissociated conformation.

Among the 12 hCFTR samples containing different protein constructs, nucleotides, and drugs for which we have generated GCERs, all but G551D-6SS-hCFTR/pATP/VX770/VX809/VX445 yield an hNBD1-dissociated structure representing a significant fraction of the population (Fig. 1E and data not shown). This protein sample, but none of the others characterized in this paper, was expressed in tissue culture cells grown in the presence of the VX809 corrector to increase yield. The compound copurified with the protein and is clearly visible in all maps (data not shown), which also clearly show the VX770 and VX445 drugs added to the purified protein (as in Fig. S3). The absence of detectable kinetically-trapped PU or hNBD1-dissociated molecules in this sample could reflect the combined action of VX809 and VX445 preventing formation of these species (Fig. 7). However, future studies will be required to test this hypothesis relative to the alternative explanation that the overall level of hNBD1-dissociated species shows some degree of sporadic variation between different hCFTR protein preparations.

## DISCUSSION

The results reported here enable formulation of a coherent model for the mechanism by which VX445 simultaneously corrects biogenesis defects in hCFTR and potentiates its chloride channel activity (Fig. 7). We present direct structural evidence that VX445 binding to a cavity formed by an N-terminal segment of the Lasso and membrane-embedded segments of TMHs 2, 10, and 11 shifts the protein population towards open-channel conformations (Figs. 1, 3 left and center, and 4A-B), explaining its potentiator activity (Fig. 7 right), and we present mutational evidence (Fig. 4C-D) that binding to the same site mediates its corrector activity (Fig. 7 center). The structure of the VX445 binding site changes significantly in conformations with and without an internal chloride channel (Fig. 3 left and center), consistent with differential binding affinity in those conformations thermodynamically driving channel opening (Fig. 7 right). VX445 binding at this TM site reverses the usual pathway of allosteric conformational control in hCFTR, pulling the NBDs together based on the conformational change in the TMDs driven by VX445, as opposed to the physiology activation pathway in which the TMDs are pulled together when ATP binding drives tight association of the NBDs. Evidence for this inversion in the direction of allosteric communication is provided by the observation that the disease-causing G551D mutation in one of the ATP interaction sites in hNBD1 prevents the NBDs from adopting their canonical ATP-bound conformation (Fig. 5) even when VX445 drives formation of the internal chloride channel in the TMDs, and it is furthermore supported by recent HDX-MS studies (Soya et al., Submitted).

**Figure 5.**
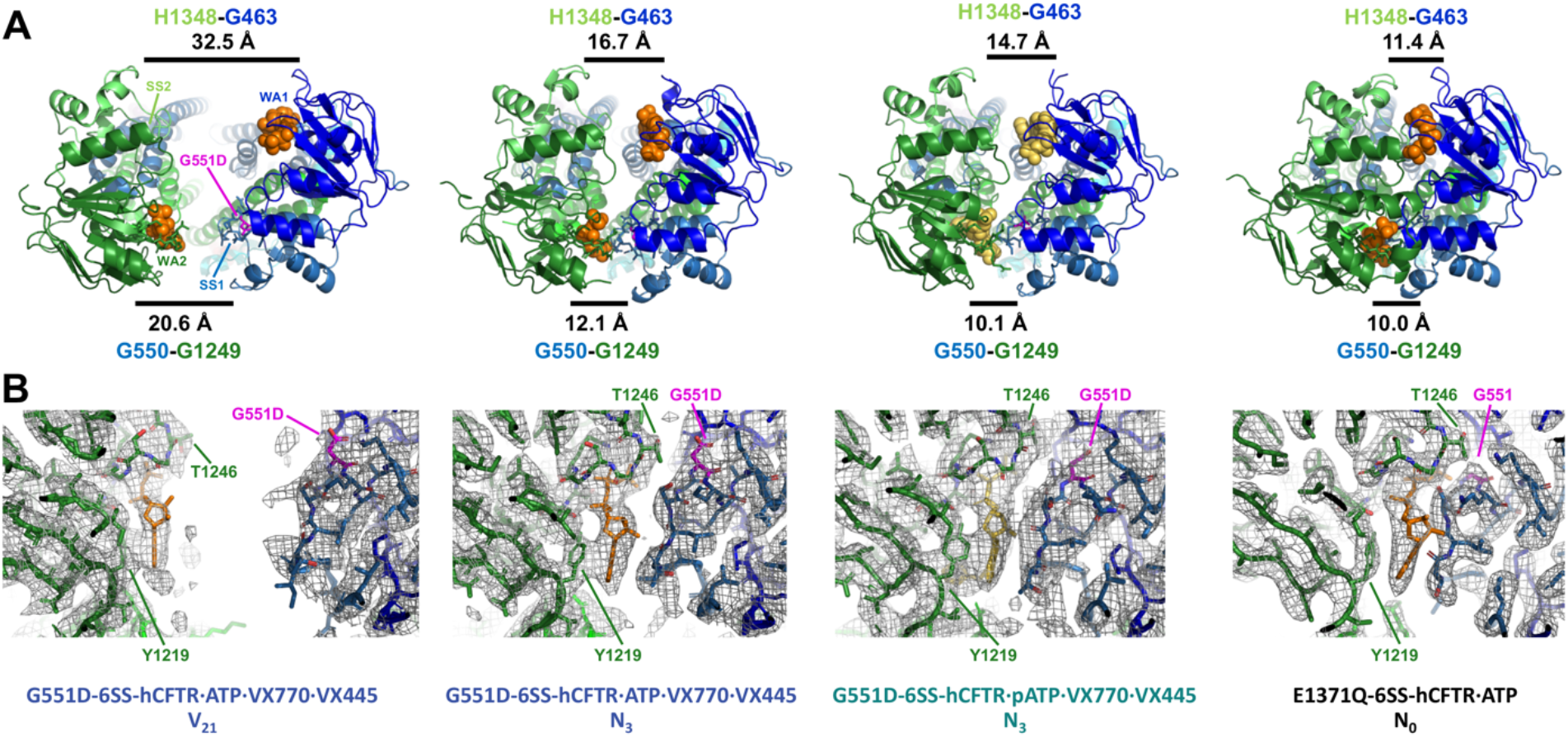
The G551D mutation prevents the NBDs in hCFTR from adopting the canonical ATP-bound structure for ABC superfamily proteins. **(A)** Top views of ribbon diagrams showing the complete NBDs from the refined models colored as in Fig. 1A. WA1 and WA2 indicate the Walker A motifs in hNBD1 and hNBD2, respectively, while SS1 and SS2 indicate the ABC superfamily Signature Sequences in the two domains. **(B)** Zoomed-in views in the same orientation of the hNBD1/2 interface in each structure at the G551D mutation site with the protein shown in stick representation colored as in panel **A** and the cryo-EM density maps displayed as a gray mesh.

The VX445 binding site driving these changes is disordered in a likely kinetically trapped, <u>p</u>artially <u>u</u>nfolded (PU) conformation of hCFTR we reconstructed at 3.4 Å (Figs. 1E and 6C-D and **Movies 4-5**). Formation of this non-native conformation *in vivo* would likely impede proper hCFTR biogenesis, particularly during the ∼45-minute period (Soya et al., Submitted) between completion of protein synthesis and export to the Golgi when nascent hCFTR is under surveillance by the folding quality-control systems in the ER (Kleizen et al., 2021; Ren et al., 2013; Soya et al., Submitted). We hypothesize that reversible dissociation of hNBD1 and the cooperatively coupled unfolding of TMH10-11 and ICL4, which mediates most packing contacts to hNBD1, gates progression into more extensively unfolded states of the TMDs (**left in** Fig. 7) and the aggregation-prone molten globule state of hNBD1 (top in Fig. 7) (Protasevich et al., 2010) during this critical post-translational period when correctors exert their major effects (Soya et al., Submitted). VX445 binding at the site identified in our structures (Figs. 3 left and **4A-B**) will strongly inhibit unfolding of TMH10-11/ICL4 (**Movies 4-5**), which will push the equilibrium towards the native hNBD1-bound conformation and away from the more extensively unfolded states likely to target nascent hCFTR for degradation. The model in Fig. 7 thus explains how VX445 binding to a single site can mediate both correction of biogenesis defects in hCFTR and potentiation of its channel activity.

**Figure 6.**
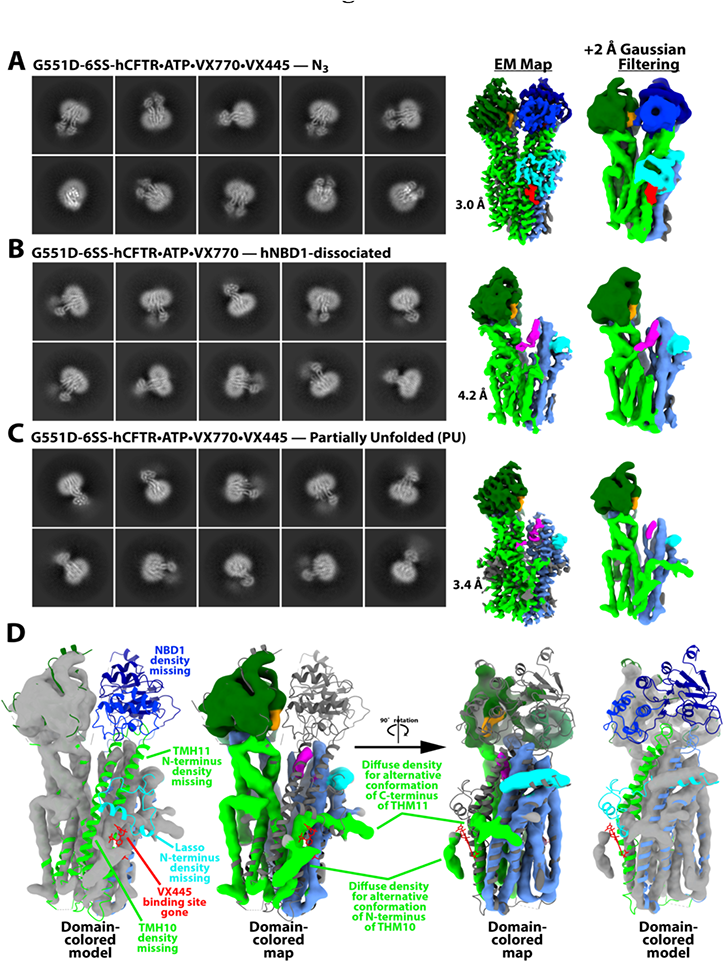
Properties of the hNBD1-dissociated and partially unfolded protein (PU) conformations generated by GCER analyses. **(A-C)** Top 10 most populated 2D classes (left) and the final non-uniform refinement map (Punjani et al., 2020) without (middle) or with 2 Å Gaussian filtering (right) for the N_3_ (panel **A**) and PU (panel **C**) structures in the G551D-6SS-hCFTR/ATP/VX770/VX445 GCER and the NBD1-dissociated structure in the G551D-6SS-hCFTR/ATP/VX770 GCER (panel **B**). Domains and ligands in the maps are colored as in Fig. 1A. **(D)** Two views of a superposition of a ribbon diagram of the N_3_ coordinate model and the Gaussian-filtered PU map using two alternative color schemes (gray or colored according to domain organization) to illustrate the unfolded regions. **Movie 4** shows a 360° rotation of the gray PU map with the ribbon diagram colored according to domain organization, while **Movie 5** shows the N3 ® PU density transition.

Recent hydrogen-deuterium exchange mass spectrometry (HDX-MS) studies (Soya et al., Submitted) show that disease-causing, destabilizing mutations in hNBD1, including the F508del mutation, consistently enhance the dynamics of TMH10-11/ICL4, which are disordered in both the PU and other hNBD1-dissociated structures we have determined (Fig. 6), while the VX809 and VX445 corrector drugs as well as stabilizing mutations in hNBD1 that suppress the effect of the F508del mutation all reverse these effects and reduce the dynamics of these protein segments. These observations together with the cryo-EM studies reported here suggest the hypothesis that dissociation of hNBD1 coupled to unfolding of TMH10-11/ICL4 represents a major obstacle to hCFTR biogenesis that contributes to the disease-causing effects of the F508del mutation (Fig. 7).

**Figure 7.**
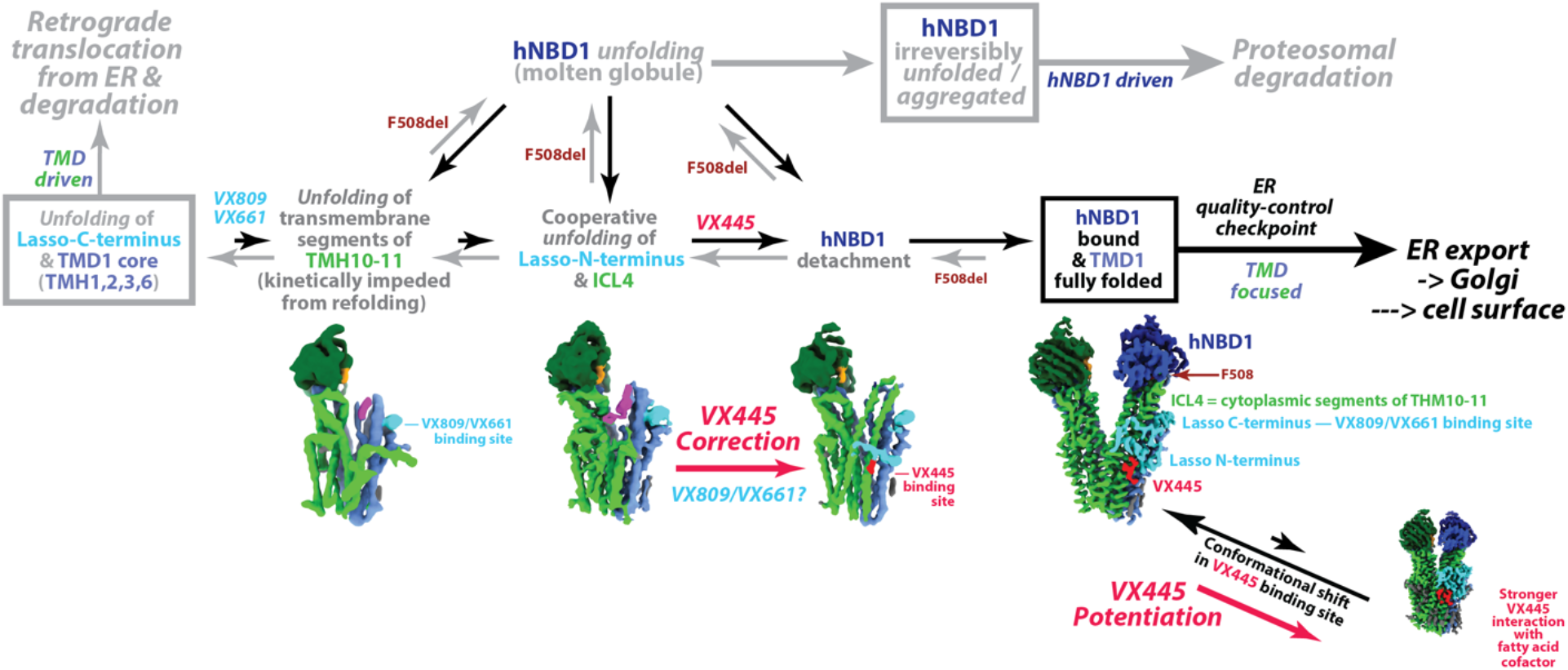
Schematic model for the mechanism of dual correction and potentiation by VX445 derived from GCER analyses of G551D-6SS-hCFTR. Protein domains and ligand are colored as in Fig. 1A. The density map with hNBD1 dissociated but ICL4 folded is one frame from **Movie 5**, calculated as described in the Methods, and could represent an average of physical states.

The structural studies reported here in cholesterol-rich digitonin micelles suggest that phospholipid interactions critically modulate hCFTR conformation and drug action (Fig. 3). Future structural studies of hCFTR should be extended to a native phospholipid bilayer environment to characterize more accurately the stereochemistry and thermodynamics of those interactions. Additional studies are also needed to evaluate the relationship of the likely kinetically-trapped, <u>p</u>artially <u>u</u>nfolded (PU) conformation of hCFTR we have reconstructed here (Fig. 6B-C) to the immaturely glycosylated, ER-resident biogenesis intermediate referred to as “Band B” in cell biological studies, and related studies will be needed to evaluate how VX809/VX661 and VX445 interact to modulate formation of this kinetically trapped species and reversibly hNBD1-dissociated species *in vivo* in the ER and in the plasma membrane as well as *in vitro* in purified hCFTR samples. Such experiments are essential to test the hypotheses (Fig. 7) that reversal of this allosteric domain-dissociation/unfolding reaction represents the mechanism-of-action of VX445 and may contribute to the mechanism-of-action VX809, breakthrough folding-corrector drugs that have effectively cured CF for patients harboring the formerly fatal F508del mutation (Keating et al., 2018; Middleton et al., 2019).

## Acknowledgements

ZY, CGB, ILU, JCK, GLL, JF, and JFH all acknowledge long-term grant support from the Cystic Fibrosis Foundation. GLL also acknowledges support from the Canadian Institute for Health Research, Cystic Fibrosis Canada, the Canada Foundation of Innovation, and NIH-NIDDK. The authors thank Elizabeth Joseloff for advice and support, and Wayne Hendrickson for helpful conversations.

## Author Contributions

Conceptualization: CW, ZY, BL, SV, DRW, RBS, CGB, ILL, JCK, GLL, JF, & JFH. Methodology: CW, ZY, BL, SV, HX, GV, OBC, FJ, RAG, WW, KHW, RBS, CGB, ILU, JCK, GLL, JF, & JFH. Investigation: CW, ZY, BL, SV, HX, GV, FJ, SS, ZR, ERM, RAG, WW, AM, ZF, KHW, JW, JCK, GLL, & JFH. Visualization: CW, ZY, BL, YL, GLL, &JFH. Funding acquisition: JCK, GLL, JF, & JFH. Project administration: JCK, GLL, JF, & JFH. Supervision: ZY, JCK, GLL, & JFH. Writing: CW, ZY, BL, OBC, DRW, CGB, ILU, JCK, GLL, JF, & JFH.

## Supplementary Information

Five movies illustrating the cryo-EM density map for the partially unfolded conformation of hCFTR and conformational transitions between and within the reconstructed conformations can be found online. The *Supplementary Information* section of this document contains a table reporting cryo-EM data collection, model refinement, and structure validation statistics followed by seven Supplementary Figures.

## Materials and data access

Expression constructs and cell lines can be obtained upon request from John C. Kappes (via e-mail to). The four cryo-EM datasets reported in Fig. 1 and Table S1 will be released in EMPAIR following acceptance of a peer-reviewed paper reporting these results. The 22 cryo-EM maps represented in Fig. 1 will be released at the same time via the EMBD along with the corresponding refined coordinate models plus three additional models refined against composite maps for the V_21_ and N_3_ conformations of G551D-6SS-hCFTR/ATP/VX770/VX445 and the N_3_ conformation of G551D-6SS-hCFTR/ATP/VX770/VX445. In the interim, the maps and models can be obtained upon request from John F. Hunt (via e-mail to).

## Methods

### Digitonin preparation

Powder from Sigma (Millipore-Sigma, St. Louis, MO) was dissolved at 10% (w/v) in purified water at 80-90 °C, aged at room temperature for 5-7 days, and then aged at 4 °C for an additional 7-21 days prior to 0.2 µM filtration and lyophilization.

### Expression and purification of full-length hCFTR

Protein constructs were expressed as previously described (Yang et al., 2018) with the addition of VX809 during cell culture for preparation of the sample used to produce the G551D-6SS-hCFTR/ATP/VX770/VX809/VX445 GCER reported in this paper (but not the other GCERs). The resulting hCFTR-expressing CHO cells were solubilized in Buffer W (20 mM HEPES, pH 7.5, 150 mM NaCl, 10 % glycerol, 0.2 mM TCEP) containing 0.5% dodecyl-β-D-maltoside, 0.1% cholesteryl hemisuccinate (CHS), 2 mM DTT, and Roche cOmplete^™^ EDTA-free Protease Inhibitor Cocktail (Millipore-Sigma, St. Louis, MO). The proteins were purified via a C-terminal STREP II tag using Strep-tactin resin (IBA LifeSciences, Göttingen, DE). The solubilizing detergents were replaced by 0.06% (w/v) digitonin via extensive washing while the protein was bound to the resin. The tagged protein was eluted in Buffer W containing 0.06% (w/v) digitonin and 4 mM *d*-desthiobiotin. The eluted protein was concentrated to ∼1 mg/ml and flash-frozen. Following thawing, the purification tags were removed by incubating with a roughly 1:1 mass ratio of TEV protease on ice for 3-5 hrs prior to Superose 6 Increase 10/300 gl (Cytivia, Marlborough, MA) gel-filtration in TMN buffer (0.06% (w/v) digitonin, 200mM NaCl, 3mM MgCl_2_, 1mM DTT, 50 mM Tris-Cl pH 7.5) with 2 mM ATP and 10% (v/v) glycerol. Fractions were flash-frozen, and hCFTR-containing fractions based on SDS-PAGE analysis were pooled after thawing and immediately phosphorylated with 1,000 U of protein kinase A (New England BioLabs) per ml for ∼1 hr on ice prior to addition of VX770 and VX445 to some samples and final purification via Superose 6 Increase 3.2/300 gl microbore gel-filtration chromatography in glycerol-free TMN buffer with 150 µM ATP. The microbore column was run on a Shimadzu (Columbia, MD) HPLC with an SPD-20A diode array detector. The resulting 100 µl fractions were flash-frozen, and hCFTR-containing samples based on SDS-PAGE analysis were pooled after thawing, supplemented with 1.85 mM ATP or 4.5 mM pATP (Axxora, Farmingdale, NY), concentrated 2-fold to 20-fold, and immediately deposited on grids. Some of the resulting concentrated stocks were flash-frozen and later thawed to make additional grids, which had very similar properties although with some reduction in the density of monomeric particles.

### VX770 preparation

A 6 µl volume of 50 mM VX770 in spectroscopic grade DMSO was added in 1 µl aliquots to 296 µl of 2% digitonin in purified water, with vigorous vortexing after each addition. The resulting solution was dialyzed in a 30 kDa Slide-A-Lyzer (Thermo Fisher, Waltham, MA) at 4 °C for 2 hours against 50 ml of purified water to remove DMSO and then centrifuged for 15 min at top speed in an Eppendorf centrifuge at 4 °C. Aliquots of the supernatant were flash-frozen, and the concentration of VX770 was calculated assuming an χ_315 nm_ of 18,900 M^-1^cm^-1^. Immediately prior to either the final glycerol-free gel-filtration purification or cryo-EM grid preparation, the drug was added to protein samples at a concentration calculated to yield a free concentration of 7 µM assuming.

### VX445 preparation

Activated charcoal (2 g) was washed three times by suspending in purified water and collecting by centrifugation prior to adding 600 mg of purified BSA (Millipore-Sigma) in 6 ml of purified water. This suspension was titrated to pH 3.5 with 2 M HCl, tumbled at 4 °C for 1 hr, titrated to pH 7.0-7.5 with 4 M NaOH, passed through a 0.22 µm filter, diluted to 10 ml final volume with purified water, and flash frozen in aliquots. Following thawing, 1 ml of this defatted BSA stock (Chen, 1967) was placed in a 1.5 ml Eppendorf tube, and 20 µl of 50 mM VX445 in spectroscopic grade DMSO was added in 1 µl aliquots with vortexing between each addition. Following centrifugation at 14,000 rpm for ∼30 minutes at 4 °C, the supernatant was removed, carefully avoiding the small black pellet, and 900 µl of it was loaded on a Superdex 200 Increase 10/300 gl gel-filtration column (Cytivia) in 150 mM NaCl, 20 mM Tris-HCl p.H 7.5. Protein-containing fractions were pooled and flash-frozen in small aliquots The concentration of VX445 was calculated assuming an χ_355 nm_ of 14,900 M^-1^cm^-1^. Immediately prior to the final glycerol-free gel-filtration purification (Fig. S3), the drug was added to protein samples to yield a final free concentration of 5 µM assuming 1:1 binding to the protein.

### Cryo-EM grid preparation

A 3 µl aliquot of freshly concentrated or thawed protein at ∼1 mg/ml was deposited on an UltraAuFoil 300 mesh R 0.6/1 grid (Quantifoil Inc., UK) at 4 °C and 100% humidity in a Vitrobot Mark IV robot (Thermo Fisher). The grid was treated in a Solarus Plasma Cleaner 950 (Gatan Inc., USA) for 25 sec with O2/H2 flow-rates of 27.5/6.4 sccm and 15 W cleaning power. Following protein deposition, grids were blotted with Whatman 1 filter paper (Whatman Inc., Piscataway, NJ) for 6-10 secs at force 3-4 prior to plunging into liquid ethane.

### Cryo-EM data collection

Cryo-EM micrographs were collected using Leginon (Cheng et al., 2021) on a 300 kV Titan Krios electron microscope (ThermoFisher Inc., USA) with a K3 camera and imaging filter (Gatan Inc., USA) at the Columbia University Cryo-EM Center using counting mode at a nominal magnification of 105,000× (nominal pixel size 0.844 Å/pixel) with a defocus range from −1.34 to −2.17 µm. Exposures of 2.5 s were dose-fractionated into 50 frames (50 ms/frame), with an exposure rate of 16 e^-^/pixel/sec for a total exposure of 56 e^-^/Å^2^.

### <u>G</u>lobal <u>C</u>onformational <u>E</u>nsemble <u>R</u>econstruction (GCER)

Image processing and 3-dimensional reconstruction were performed using cryoSPARC 3.2 (Punjani and Fleet, 2021; Punjani et al., 2020), yielding the results given in Table S1 and illustrated in Fig. S4. GCER was performed using the previously described iterative heterorefinement procedure (Li et al., 2022), which is summarized in the main text, after an exhaustive search for alternative conformations in preliminary structures that included minimally 3D-VA at 4 Å using the full molecular envelope, 2-3 class *ab initio* reconstruction at 8-4 Å, and sometimes additionally heteroclassification using duplicate volumes or 3D classification with a local mask in conformationally variable regions identified via 3D-VA or in previous samples.

### Coordinate refinement

Manual rebuilding was performed using COOT (Casanal et al., 2020), and real-space coordinate refinement and model-validation calculations were performed in PHENIX (Afonine et al., 2018). After an initial round of refinement against a new cryo-EM map including ADP refinement, all B-factors in the model were reset to the average value followed by one round of refinement without ADP refinement and then a second round including ADP refinement followed by additional rounds of manual rebuilding and computational refinement if needed.

### Movies

Movies show 7-frame (Movie 5) or 15-frame (Movies 1-3) intermediates displays of 3D-VAs performed in cryoSPARC using particle orientations from a homogeneous refinement. This input refinement was performed on particle stacks from the final cycle of iterative heteroclassification (Figs. 1 and S4) seeded with the volume from non-uniform refinement for analysis of a single conformational class or the volume from a 1-class ab initio reconstruction for analysis of combined conformational classes. The envelope for 3D-VA was produced by aligning the refined coordinate model for the class, or both coordinate models superimposed for combined classes, with the input map, using molmap in Chimera with a 3.5 Å radius to convert the model(s) to a map, and thresholding that map at density level of 0.001, filling holes, dilating by 2 Å, and adding a 15 voxel soft pad in cryoSPARC. Each movie frame shows a weighted average of particles in 3 adjacent eigenvalue bins from one eigenmode in the covariance matrix between all pairs of points in the map from homogeneous refinement. Note that these movies may not represent physically accurate stereochemical trajectories (Seitz and Frank, 2020), but they are calculated directly from subsets of particles in the dataset, so all frames represent an average of conformations present in the protein sample. Reduction in the volume of the cryo-EM density in some regions in some frames likely reflects disorder either in the form of rigid-body movement (rotational/translational diffusion) of natively folded (sub)domains or 2° structure elements or potentially local backbone unfolding.

### Molecular graphics

Images showing surface renderings of cryo-EM maps and movies were prepared using ChimeraX (Pettersen et al., 2021), while all other molecular images including mesh representations of maps were prepared using PyMOL (http://www.pymol.org/) (Chen, 1967).

### Cell lines and antibodies

Generation of CFBE41o-cells lines (a gift from D. Gruenert, University of California, San Francisco) expressing inducible hCFTR variants with a 3HA-tag in the fourth extracellular loop was described previously (Veit et al., 2012). Cells were grown in MEM medium containing 10% fetal bovine serum, 10 mM 4-(2-hydroxyethyl)-1-piperazineethanesulfonic acid (HEPES) and 2 mM L-glutamine, and hCFTR expression was induced for ≥3 days with 250-500 ng/ml doxycycline. New hCFTR variants were generated by overlapping PCR mutagenesis (Du et al., 2005). Monoclonal mouse anti-CFTR 660 Ab to hNBD1) was provided by J. Riordan and the CF Foundation via the CFTR Antibodies Distribution Program. Immunoblotting with anti-Na^+^/K^+^-ATPase Ab SC-48345 (Santa Cruz Biotechnology, Dallas, TX) was used as a loading control.

### Quantitative immunoblotting

Equal amounts of CFBE14o-lysates expressing CFTR-3HA variants were separated by SDS-PAGE, blotted to nitrocellulose, and probed with a monoclonal anti-HA primary Ab as described (*102*). Digital images of enhanced chemiluminescence signals (ECL-Pico, ThermoFisher) were collected from immunoblots using the BioRad Chemidoc MP Imaging System and, after background correction, were quantified using the Image Lab 6.1 (BioRad) and NIH Image 2.0 software (http://rsb.info.nih.gov/nih-image/).

### Short-circuit current measurements

I_sc_ measurement were performed essentially as described (*100*). G551D-, G551D+M1105I-, or G551D+I1109F-CFTR-expressing CFBE14o-cells were polarized on Snapwell filters and mounted in Ussing chambers (Physiologic Instruments, Reno, NV) in Krebs-bicarbonate Ringer buffer (140 mM Na^+^, 120 mM Cl^−^, 5.2 mM K^+^, 25 mM HCO_3_^−^, 2.4 mM HPO_4_, 0.4 mM H_2_PO_4_, 1.2 mM Ca^2+^, 1.2 mM Mg^2+^, 5 mM glucose, pH 7.4), which was mixed by bubbling with carbogen (95% O_2_ and 5% CO_2_). To generate a basolateral-to-apical chloride gradient, NaCl was replaced with 115 mM Na^+^ gluconate in the apical buffer, the basolateral membrane was permeabilized with 100 µM amphotericin B. I_sc_ was determined in the presence of 100 μM amiloride. Transepithelial voltage was clamped at 0 mV (VCC MC8 Multichannel Voltage/current Clamp, Physiologic Instruments) after compensating for voltage offsets. Current and resistance were recorded at 37°C with the Acquire and Analyze package (Physiologic Instruments). Following 20 µM forskolin-induced cAMP-dependent PKA activation and 3 µM VX-770 potentiation of the channel, VX-445 dose-response was recorded.

## Supplementary Information

### Movies

**Movie 1 (associated with** Fig. 1**): *Major protein conformational changes leading to internal chloride channel formation in the GCER for G551D-6SS-hCFTR/ATP/VX770/VX445.*** Sequences of cryo-EM density maps colored according to domain organization (as in Fig. 1A) calculated from 3D-VA at 5 Å resolution of combined particles in the V_21_ and V_17_ (left) or V_17_ and N_3_ (right) conformational classes after the final round of iterative heterorefinement (Fig. S4). Both movies represent eigenmode 0 from 3D-VA performed as described in the Methods.

**Movie 2 (associated with** Fig. 1**): *Residual conformational fluctuations in the V_21_ conformational class of G551D-6SS-hCFTR/ATP/VX770/VX445.*** Sequence of cryo-EM density maps colored according to domain organization (as in Fig. 1A) calculated from 3D-VA at 5 Å resolution of the particles in the V_21_ conformational class after the final round of iterative heterorefinement (Fig. S4). The movies represent eigenmodes 0 and 6 from 3D-VA performed as described in the Methods.

**Movie 3 (associated with** Fig. 1**): *Residual conformational fluctuations in the N_3_ conformational class of G551D-6SS-hCFTR/ATP/VX770/VX445.*** Sequence of cryo-EM density maps colored according to domain organization (as in Fig. 1A) calculated from 3D-VA at 5 Å resolution of the particles in the N_3_ conformational class after the final round of iterative heterorefinement (Fig. S4). The movies represent eigenmodes 0 and 4 from 3D-VA performed as described in the Methods.

**Movie 4 (associated with** Fig. 6**): *Cryo-EM map for the <u>p</u>artially <u>u</u>nfolded (PU) conformational class of G551D-6SS-hCFTR/ATP/VX770/VX445.*** The map from the final round of iterative heterorefinement (Fig. S4) colored according to domain organization (as in Fig. 1A) is shown on the left, while the same map is shown on the right in gray semi-transparent surface representation superimposed on a ribbon diagram of the natively folded N_3_ conformation reconstructed from the same protein sample.

**Movie 5 (associated with** Fig. 6**): *Cryo-EM density transition between the N_3_ and <u>p</u>artially <u>u</u>nfolded (PU) conformational classes.*** Sequence of maps colored according to domain organization (as in Fig. 1A) calculated from 3D-VA at 5 Å resolution of combined particles in the N_3_ and PU classes after the final round of iterative heterorefinement (Fig. S4). The movie represents eigenmode 0 from 3D-VA performed as described in the Methods.

---

**Supplementary Table S1 (associated with Fig. 1):** Cryo-EM data collection, refinement and validation statistics.

| Protein construct / ligands | G551D-6SS-hCFTR/ATP/VX770/VX445 |  |  |  | E1371Q-6SS-hCFTR/ATP |
| --- | --- | --- | --- | --- | --- |
| TMD opening angle (°) | 2.7 | 17.6 | 21.9 | -3.8 | 0 |
| hNBD1 Core rotation (°) | 1.7 | 5.5 | 9.3 | N/A | 0 |
| Notes | N<sub>3</sub> | V<sub>17</sub> | V<sub>21</sub> | Partially Unfolded – PU<br>(NBD1-dissociated) | N<sub>0</sub><br>Reference |
| PDB deposition code |  |  |  |  |  |
| EMDB deposition code |  |  |  |  |  |
| **Data collection & processing** |  |  |  |  |  |
| EMPIAR deposition code |  |  |  |  |  |
| Microscope | Columbia Krios C1 |  |  |  | Columbia C1 |
| Nominal magnification | 105,000 |  |  |  | 105,000 |
| Voltage (kV) | 300 |  |  |  | 300 |
| Electron exposure (e<sup>-</sup>/Å<sup>2</sup>) | 57.10 |  |  |  | 57.10 |
| Defocus range (μm) | -0.45/-2.5 |  |  |  | -0.66/-2.9 |
| Pixel size (Å) | 0.837 |  |  |  | 0.837 |
| Micrographs used (#) | 13,304 |  |  |  | 6861 |
| Total particles picked (#) | 2,432,657 |  |  |  | 1,135,813 |
| Particles after 2D cleaning (#) | 1,118,124 |  |  |  | 655,958 |
| Final particles assigned (#) | 703,530 |  |  |  | 380383 |
| Percentage final/2D (%) | 62.9 |  |  |  | 58.1 |
| Relative percentage (%) | 25.4 | 21.6 | 38.7 | 14.3 | 26.9 |
| Symmetry imposed | C1 | C1 | C1 | C1 | C1 |
| Map resolution (Å) | 3.0 | 3.4 | 2.9 | 3.5 | 3.0 |
| FSC threshold | 0.143 | 0.143 | 0.143 | 0.143 | 0.143 |
| **Refinement** |  |  |  |  |  |
| Initial model used (PDB ID) |  |  |  |  | 6MSM |
| Model resolution (Å) | 3.0/3.3 | 3.5/3.8 | 2.8/3.0 | 3.4/3.7 | 3.0/3.2 |
| FSC threshold | 0.143/0.5 | 0.143/0.5 | 0.143/0.5 | 0.143/0.5 | 0.143/0.5 |
| Map sharpening *B* factor (Å<sup>2</sup>) | -75.4 | -85.2 | -76.0 | -73.5 | -77.9 |
| **Model composition** |  |  |  |  |  |
| Non-hydrogen atoms | 10378 | 9677 | 9791 | 7158 | 10092 |
| Protein residues | 1155 | 1147 | 1143 | 887 | 1166 |
| Ligands | 48 | 28 | 31 | 9 | 42 |
| ***B* factors (Å<sup>2</sup>)** |  |  |  |  |  |
| Protein | 100.11 | 107.65 | 99.13 | 125.14 | 60.96 |
| Ligand | 115.74 | 91.35 | 113.40 | 90.68 | 67.26 |
| **R.m.s. deviations** |  |  |  |  |  |
| Bond lengths (Å) | 0.003 (0) | 0.003 (0) | 0.003 (0) | 0.003 (0) | 0.004 |
| Bond angles (°) | 0.620 (1) | 0.665 (0) | 0.520 (0) | 0.670 (1) | 0.59 |
| **Validation** |  |  |  |  |  |
| MolProbity score | 1.86 | 2.04 | 1.57 | 1.95 | 1.52 |
| Clashscore | 15.36 | 13.63 | 7.52 | 13.41 | 8.3 |
| Poor rotamers (%) | 0.10 | 0.10 | 0.00 | 0.00 | 0.00 |
| **Ramachandran plot** |  |  |  |  |  |
| Favored (%) | 97.03 | 94.09 | 97.09 | 95.51 | 97.67 |
| Allowed (%) | 2.97 | 5.56 | 2.74 | 4.49 | 2.33 |
| Disallowed (%) | 0.00 | 0.35 | 0.18 | 0.00 | 0.00 |

**Supplementary Table S1 (associated with Fig. 1): Cryo-EM data collection, refinement and validation statistics (continued).**
| Protein construct / ligands | G551D-6SS-hCFTR/pATP/VX770/VX809/VX445 |  |  |
| --- | --- | --- | --- |
| TMD opening angle (°) | 2.7 | 18.8 | 21.4 |
| hNBD1 Core rotation (°) | 2.5 | 5.9 | 9.9 |
| Notes | N<sub>3</sub> | V<sub>17</sub> | V<sub>21</sub> |
| PDB deposition code |  |  |  |
| EMDB deposition code |  |  |  |
| **Data collection & processing** |  |  |  |
| EMPIAR deposition code |  |  |  |
| Microscope |  | Columbia Krios C1 |  |
| Nominal magnification |  | 105,000 |  |
| Voltage (kV) |  | 300 |  |
| Electron exposure (e<sup>-</sup>/Å<sup>2</sup>) |  | 57.10 |  |
| Defocus range (μm) |  | -0.70/-2.8 |  |
| Pixel size (Å) |  | 0.837 |  |
| Micrographs used (#) |  | 7341 |  |
| Total particles picked (#) |  | 813155 |  |
| Particles after 2D cleaning (#) |  | 344357 |  |
| Final particles assigned (#) |  | 155150 |  |
| Percentage final/2D (%) |  | 45.1 |  |
| Relative percentage (%) | 32.6 | 20.9 | 46.5 |
| Symmetry imposed | C1 | C1 | C1 |
| Map resolution (Å) | 3.2 | 3.9 | 3.2 |
| FSC threshold | 0.143 | 0.143 | 0.143 |
| **Refinement** |  |  |  |
| Initial model used (PDB ID) |  |  |  |
| Model resolution (Å) | 3.1/3.6 | 3.8/6.8 | 3.1/3.6 |
| FSC threshold | 0.143/0.5 | 0.143/0.5 | 0.143/0.5 |
| Map sharpening *B* factor (Å<sup>2</sup>) | -50.0 | -57.1 | -53.2 |
| Model composition |  |  |  |
| Non-hydrogen atoms | 9928 | 9845 | 9806 |
| Protein residues | 1167 | 1142 | 1148 |
| Ligands | 29 | 40 | 25 |
| *B* factors (Å<sup>2</sup>) |  |  |  |
| Protein | 129.37 | 16.82 | 112.73 |
| Ligand | 104.86 | 158.44 | 89.23 |
| R.m.s. deviations |  |  |  |
| Bond lengths (Å) | 0.003 | 0.10 | 0.003 |
| Bond angles (°) | 0.678 | 1.097 | 0.635 |
| Validation |  |  |  |
| MolProbity score | 1.84 | 2.27 | 1.87 |
| Clashscore | 13.17 | 23.01 | 12.07 |
| Poor rotamers (%) | 0 | 0.1 | 0.1 |
| Ramachandran plot |  |  |  |
| Favored (%) | 96.72 | 93.63 | 96.05 |
| Allowed (%) | 3.28 | 6.11 | 3.95 |
| Disallowed (%) | 0.00 | 0.27 | 0.00 |

Supplementary Table S1 (associated with Fig. 1): Cryo-EM data collection, refinement and validation statistics (continued).
| Protein construct / ligands | G551D-6SS-hCFTR/ATP/VX770 |  |  |  |  |  |  |
| --- | --- | --- | --- | --- | --- | --- | --- |
| TMD opening angle (°) | 3.8 | 15.5 | 20.0 | 21.2 | 23.78 | 30.8 | 22.1 |
| hNBD1 Core rotation (°) | N/A | 6.8 | 9.8 | 8.9 | 8.9 | 8.9 | 7.3 |
| Notes | NBD1-dissociated | V<sub>17</sub> | V<sub>21</sub> | V<sub>21</sub>LR<br>Lipid raft | V<sub>23</sub> | V<sub>31</sub> | Weak NBD2 |
| PDB deposition code |  |  |  |  |  |  |  |
| EMDB deposition code |  |  |  |  |  |  |  |
| Data collection & processing |  |  |  |  |  |  |  |
| EMPIAR deposition code |  |  |  |  |  |  |  |
| Microscope | Columbia Krios C1 |  |  |  |  |  |  |
| Nominal magnification | 105,000 |  |  |  |  |  |  |
| Voltage (kV) | 300 |  |  |  |  |  |  |
| Electron exposure (e<sup>-</sup>/Å<sup>2</sup>) | 57.10 |  |  |  |  |  |  |
| Defocus range (μm) | -0.52/-2.5 |  |  |  |  |  |  |
| Pixel size (Å) | 0.837 |  |  |  |  |  |  |
| Micrographs used (#) | 10255 |  |  |  |  |  |  |
| Total particles picked (#) | 2,242,326 |  |  |  |  |  |  |
| Particles after 2D cleaning (#) | 657,837 |  |  |  |  |  |  |
| Final particles assigned (#) | 525,497 |  |  |  |  |  |  |
| Percentage final/2D (%) | 80.0 |  |  |  |  |  |  |
| Relative percentage (%) | 10.4 | 15.1 | 17.2 | 18.2 | 16.5 | 13.1 | 9.5 |
| Symmetry imposed | C1 | C1 | C1 | C1 | C1 | C1 | C1 |
| Map resolution (Å) | 4.0 | 3.9 | 3.5 | 3.6 | 3.5 | 4.2 | 5.8 |
| FSC threshold | 0.143 | 0.143 | 0.143 | 0.143 | 0.143 | 0.143 | 0.143 |
| Refinement |  |  |  |  |  |  |  |
| Initial model used (PDB ID) |  |  |  |  |  |  |  |
| Model resolution (Å) | 4.1/6.4 | 3.8/4.2 | 3.4/3.7 | 3.5/3.8 | 3.8/6.4 | 4.1/6.1 | 5.9/7.5 |
| FSC threshold | 0.143/0.5 | 0.143/0.5 | 0.143/0.5 | 0.143/0.5 | 0.143/0.5 | 0.143/0.5 | 0.143/0.5 |
| Map sharpening *B* factor (Å<sup>2</sup>) |  |  |  |  |  |  |  |
| Model composition |  |  |  |  |  |  |  |
| Non-hydrogen atoms | 7130 | 9357 | 9569 | 10435 | 8922 | 8889 | 9222 |
| Protein residues | 885 | 1121 | 1137 | 1148 | 1103 | 1104 | 1142 |
| Ligands | 2 | 19 | 23 | 46 | 3 | 3 | 4 |
| *B* factors (Å<sup>2</sup>) |  |  |  |  |  |  |  |
| Protein | 212.10 | 192.31 | 165.66 | 92.78 | 135.65 | 194.05 | 179 |
| Ligand | 188.66 | 174.63 | 151.26 | 101.10 | 168.84 | 221.19 | 268 |
| R.m.s. deviations |  |  |  |  |  |  |  |
| Bond lengths (Å) | 0.003 (0) | 0.003 (0) | 0.003 (0) | 0.003 (0) | 0.003 (0) | 0.003 (0) | 0.003(0) |
| Bond angles (°) | 0.684 (5) | 0.760 (6) | 0.594 (1) | 0.648 (1) | 0.774 (14) | 0.758 (8) | 0.913(12) |
| Validation |  |  |  |  |  |  |  |
| MolProbity score | 2.12 | 2.16 | 1.93 | 2.14 | 2.26 | 2.21 | 3.98 |
| Clashscore | 15.53 | 19.87 | 14.40 | 27.25 | 17.03 | 19.44 | 24.85 |
| Poor rotamers (%) | 0.00 | 0.10 | 0.00 | 0.00 | 0.00 | 0.00 | 0.2 |
| Ramachandran plot |  |  |  |  |  |  |  |
| Favored (%) | 93.47 | 94.69 | 96.10 | 96.58 | 90.74 | 93.50 | 95.85 |
| Allowed (%) | 6.53 | 5.04 | 3.90 | 3.42 | 8.98 | 5.86 | 3.98 |
| Disallowed (%) | 0.00 | 0.27 | 0.00 | 0.00 | 0.27 | 0.64 | 0.18 |

**Supplementary Table S1 (associated with Fig. 1): Cryo-EM data collection, refinement and validation statistics (concluded).**
| Supplementary Table S1 (associated with Fig. 1). Cryo-EM data collection, refinement and validation statistics (continued). |  |  |  |  |  |  |  |
| --- | --- | --- | --- | --- | --- | --- | --- |
| Protein construct / ligands | G551D-6SS-hCFTR/ATP *apo* |  |  |  |  |  |  |
| TMD opening angle (°) | 15.0 | 20.8 | 21.0 | 23.8 | 23.8 | 31.0 | 22.1 |
| hNBD1 Core rotation (°) | 5.5 | 9.8 | 8.8 | 9.1 | 8.5 | 8.9 | 7.3 |
| Notes | NBD1-dissociated | V<sub>17</sub> | V<sub>21</sub> | V<sub>21</sub>LR<br>Lipid raft | V<sub>23</sub> | V<sub>31</sub> | Weak<br>NBD2 |
| PDB deposition code |  |  |  |  |  |  |  |
| EMDB deposition code |  |  |  |  |  |  |  |
| **Data collection & processing** |  |  |  |  |  |  |  |
| EMPIAR deposition code |  |  |  |  |  |  |  |
| Microscope | Columbia Krios C1 |  |  |  |  |  |  |
| Nominal magnification | 105,000 |  |  |  |  |  |  |
| Voltage (kV) | 300 |  |  |  |  |  |  |
| Electron exposure (e<sup>-</sup>/Å<sup>2</sup>) | 57.10 |  |  |  |  |  |  |
| Defocus range (μm) | -0.70/-2.8 |  |  |  |  |  |  |
| Pixel size (Å) | 0.837 |  |  |  |  |  |  |
| Micrographs used (#) | 6565 |  |  |  |  |  |  |
| Total particles picked (#) | 678,685 |  |  |  |  |  |  |
| Particles after 2D cleaning (#) | 493,476 |  |  |  |  |  |  |
| Final particles assigned (#) | 252,682 |  |  |  |  |  |  |
| Percentage final/2D (%) | 51.2 |  |  |  |  |  |  |
| Relative percentage (%) | 9.7 | 15.7 | 19.1 | 17.0 | 15.1 | 9.5 | 13.8 |
| Symmetry imposed | C1 | C1 | C1 | C1 | C1 | C1 | C1 |
| Map resolution (Å) | 6.8 | 4.4 | 3.7 | 3.9 | 4.5 | 7.1 | 5.3 |
| FSC threshold | 0.143 | 0.143 | 0.143 | 0.143 | 0.143 | 0.143 | 0.143 |
| **Refinement** |  |  |  |  |  |  |  |
| Initial model used (PDB ID) |  |  |  |  |  |  |  |
| Model resolution (Å) | 6.5/7.6 | 4.4/6.8 | 3.7/4.2 | 3.9/4.3 | 4.5/6.8 | 6.9/7.7 | 5.3/7.0 |
| FSC threshold | 0.143/0.5 | 0.143/0.5 | 0.143/0.5 | 0.143/0.5 | 0.143/0.5 | 0.143/0.5 | 0.143/0.5 |
| Map sharpening *B* factor (Å<sup>2</sup>) | -566.2 | -117.4 | -68.2 | -78.2 | -125.0 | -362.7 | -321.0 |
| Model composition |  |  |  |  |  |  |  |
| Non-hydrogen atoms | 7130 | 9357 | 9423 | 10440 | 8922 | 8889 | 9222 |
| Protein residues | 885 | 1121 | 1135 | 1148 | 1103 | 1104 | 1142 |
| Ligands | 2 | 19 | 18 | 45 | 3 | 3 | 4 |
| *B* factors (Å<sup>2</sup>) |  |  |  |  |  |  |  |
| Protein | 98.75 | 294.06 | 208.54 | 188.94 | 262.05 | 194.05 | 268.48 |
| Ligand | 79.38 | 274.24 | 197.39 | 244.65 | 281.11 | 221.19 | 337.58 |
| R.m.s. deviations |  |  |  |  |  |  |  |
| Bond lengths (Å) | 0.003(0) | 0.003 (0) | 0.003 (0) | 0.003 (0) | 0.024 (7) | 0.003(0) | 0.005 (0) |
| Bond angles (°) | 0.770(2) | 0.894 (11) | 0.691 (1) | 0.702 (6) | 0.844 (11) | 0.817(3) | 0.937 (16) |
| Validation |  |  |  |  |  |  |  |
| MolProbity score | 2.01 | 2.23 | 2.07 | 2.22 | 2.38 | 2.22 | 2.37 |
| Clashscore | 14 | 24.31 | 18.40 | 31.04 | 24.11 | 19.55 | 34.21 |
| Poor rotamers (%) | 0 | 0.00 | 0.00 | 0.10 | 0.00 | 0 | 0.20 |
| Ramachandran plot |  |  |  |  |  |  |  |
| Favored (%) | 95.07 | 94.87 | 95.65 | 96.40 | 91.38 | 93.32 | 94.88 |
| Allowed (%) | 4.93 | 4.77 | 4.35 | 3.60 | 8.25 | 6.23 | 4.95 |
| Disallowed (%) | 0 | 0.36 | 0.00 | 0.00 | 0.37 | 0.46 | 0.18 |

**Figure S1 (associated with Fig. 1).**
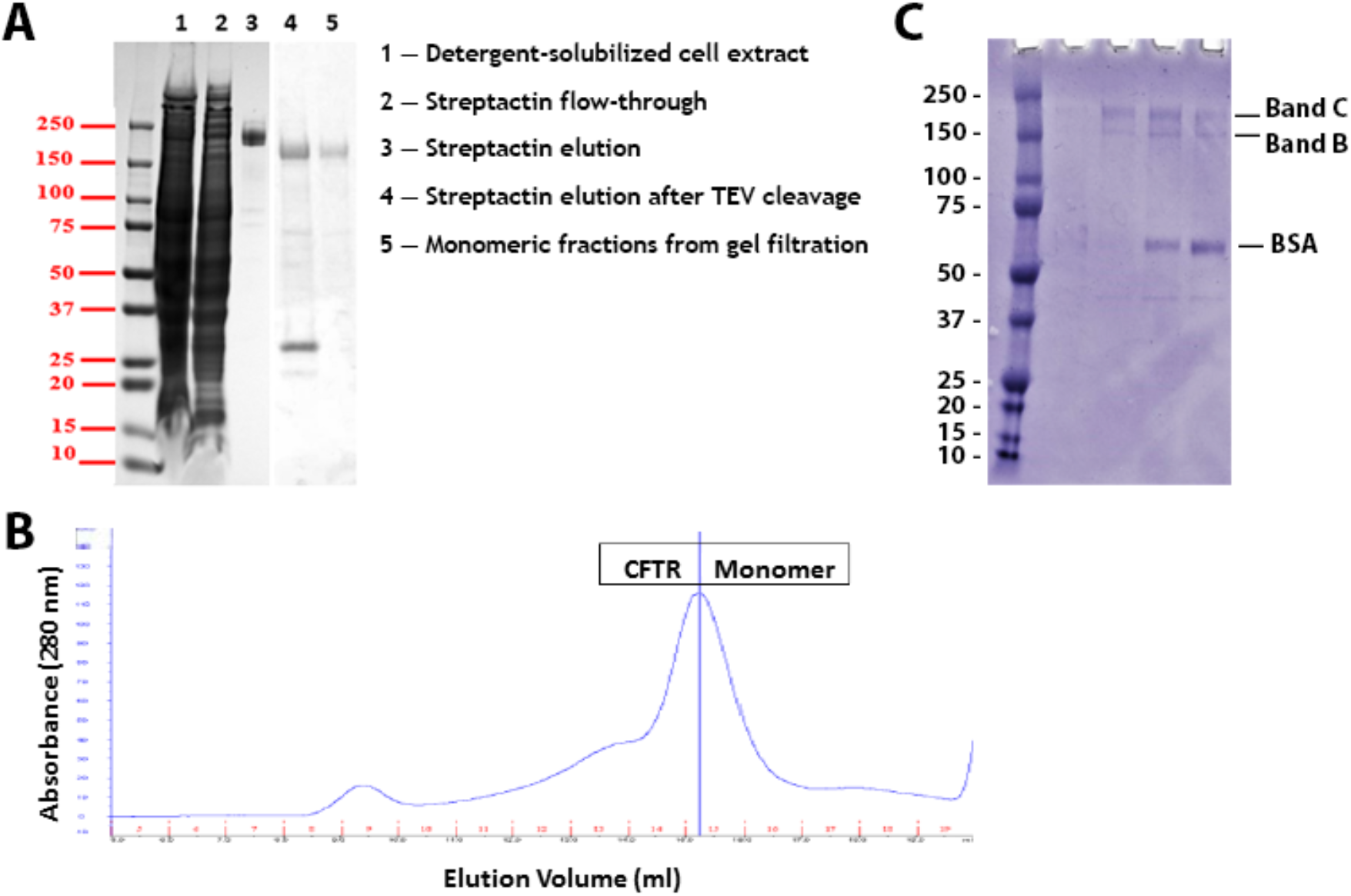
Representative hCFTR purification data. **(A)** Coomassie-Blue-stained SDS-PAGE gel showing progress of a representative purification from extraction from CHO cells through the first gel-filtration column in glycerol-containing buffer. **(B)** Representative trace from the first gel-filtration purification step on Superose 6 Increase 10/300 GL column in glycerol-containing buffer monitored by absorbance at 280 nm. **(C)** Coomassie-Blue-stained SDS-PAGE gel showing fractions from the second gel-filtration chromatography purification step on a microbore Superose 6 Increase 3.2/300 GL gel-filtration column running in glycerol-free buffer. This gel shows the protein used to produce the samples for the G551D-6SS-hCFTR/ATP/VX770/VX445 GCER shown in Figs. 1 and 3-6. VX770 and BSA•VX445 were added to this protein sample shortly before microbore gel-filtration chromatography. The two fractions with the highest protein concentrations based on Coomassie staining in the gel in panel **C** were pooled, brought to 2 mM ATP, concentrated ∼8-fold, and deposited on cryo-EM grids within 4 hours of completion of the column run without being frozen after elution from the last column. Gel-filtration was run at 4 °C in TMN buffer (0.06% (w/v) digitonin, 200mM NaCl, 3mM MgCl_2_, 1mM DTT, 50 mM Tris-Cl pH 7.5) with 2 mM ATP and 10% (v/v) glycerol for the first glycerol-containing column and 150 µM ATP for the second glycerol-free column. See Methods for additional information.

**Figure S2 (associated with Fig. 1).**
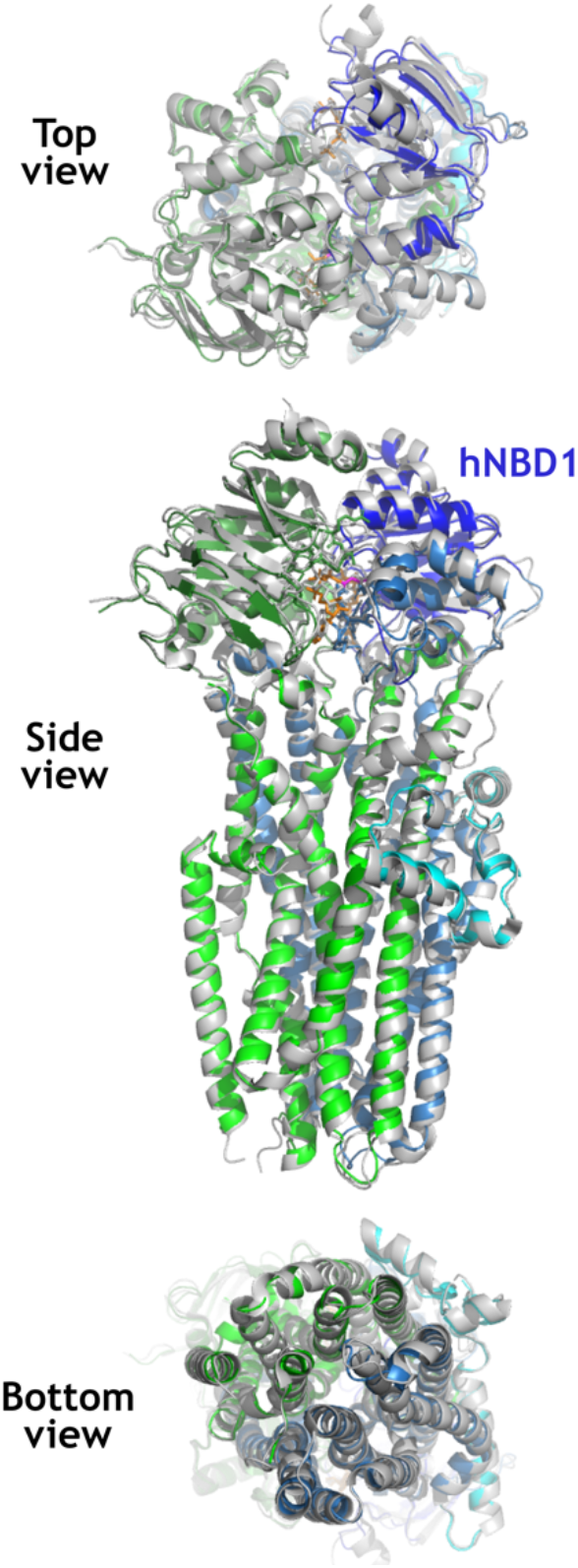
The 6SS mutation set that thermodynamically stabilizes hNBD1 does not significantly alter the structure of hCFTR. The ribbon diagrams show least-squares alignment of a 3.0 Å cryo-EM structure of the E1371Q-6SS-hCFTR base construct employed in the studies reported here (colored as in Fig. 1A) and the previously reported 3.2 Å cryo-EM structure of E1371Q-hCFTR without any mutations other than E1371Q. This mutation prevents ATP hydrolysis at the Walker A site in hNBD2, and both structures show clear electron density for unhydrolyzed ATP at both ATP-binding sites. The Walker A site in hNBD1 is catalytically inactive in native hCFTR, presumably due to sequence divergence in the Walker B sequence in hNBD1 and the Signature Sequence of hNBD2, which form one of the two composite ATP-binding sites at the interface between hNBD1-hNBD2. The root-mean-square-deviation (RMSD) is 0.75 Å for alignment of 1097 Cα atoms from 1166 residues present in both models. The apparent conformational change in the exit pore region, which is > 70 Å away from the nearest 6SS mutation site, reflects a difference in interpretation of two maps that look equivalent in that region.

**Figure S3 (associated with Fig. 1).**
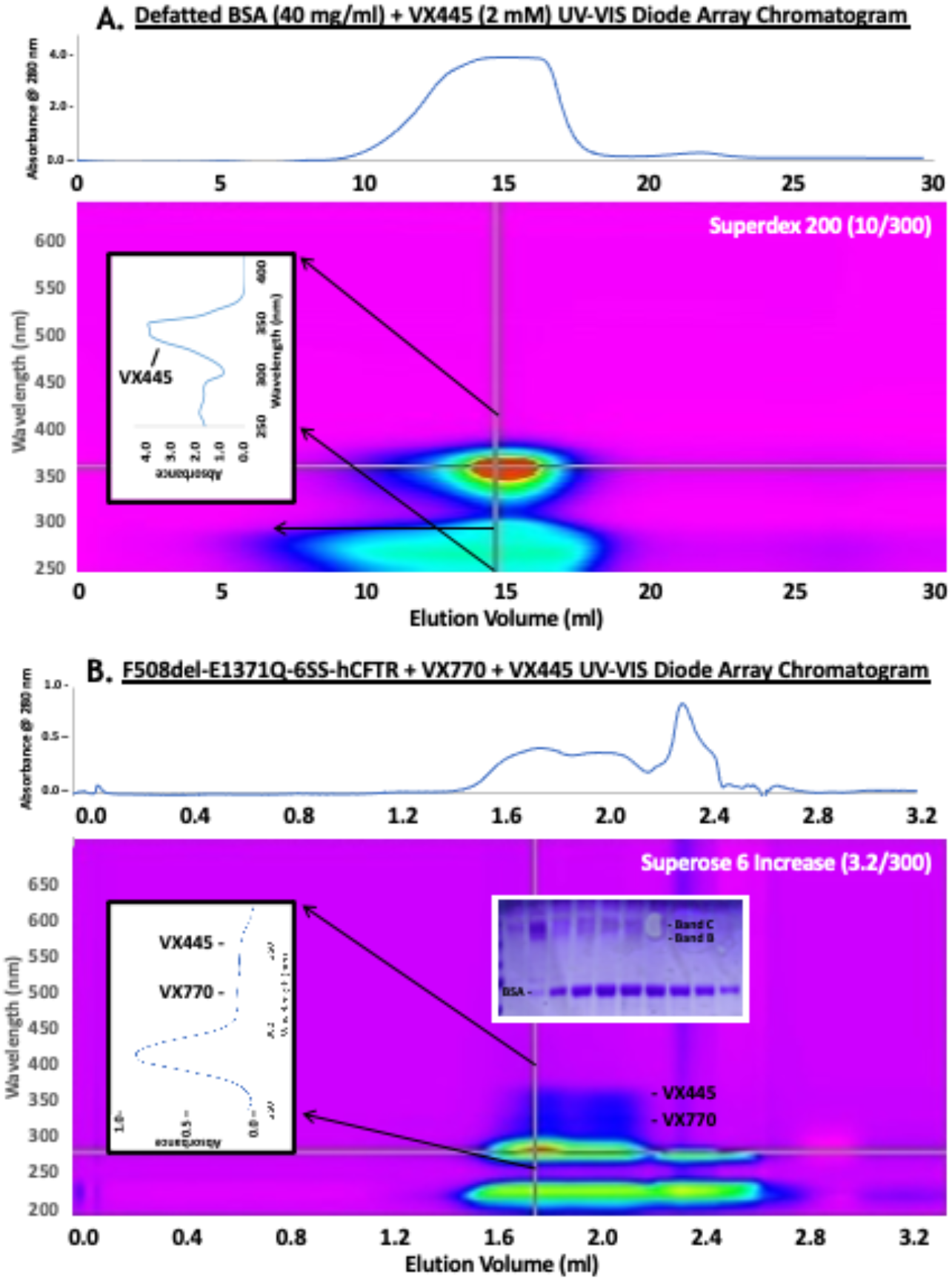
BSA host-guest-like method to deliver VX445 to hCFTR without drug aggregation. (**A)** VX445 at 50 mM in DMSO was diluted to 2 mM in 40 mg/ml defatted BSA prior to gel filtration in a simple detergent-free buffer. (**B)** Microbore gel-filtration chromatography of purified G551D-6SS-hCFTR mixed with an aliquot of the peak fraction from panel **A** and also VX770 solubilized at ∼600 µM in 2% (w/v) digitonin. The spectrum of the peak hCFTR fraction (left inset) nominally indicates a 2:2:1 ratio of VX445:VX770:hCFTR, but the 150 µM ATP in the buffer in this column distorts the absorbance at 280 nm, making this estimate an upper-limit for the drug-protein ratio. The column and buffer are described the legend for Fig. S1C. See Methods for more information.

**Figure. S4 (associated with Fig. 1):**
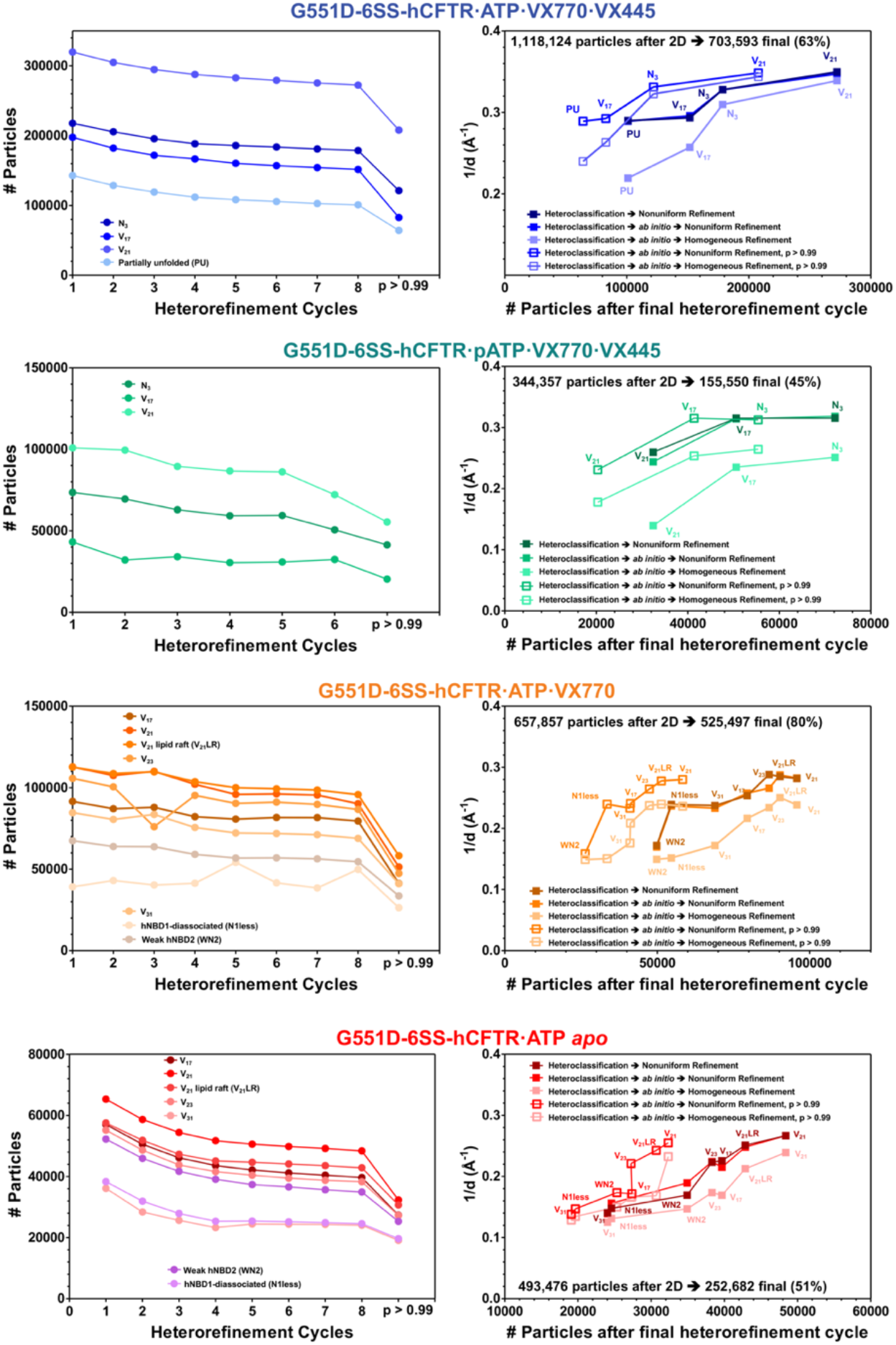
Conformational class assignments during iterative heterorefinement and validation analyses on the final classes. The p ≥ 0.99 label indicates reconstructions from the subset of particles assigned to each class that have at least a 99% probability of belonging to that class in the final heterorefinement cycle in cryoSPARC (Punjani and Fleet, 2021; Punjani et al., 2020). Classes are labeled with the abbreviations in Fig. 1E except for V_21_ lipid raft (V_21_LR), hNBD1-dissociated (N1less) and weak hNBD2 density (WN2). Volume classes with significantly lower resolution in homogeneous refinement compared to non-uniform refinement have more heterogeneity among their particles, reflecting either higher conformational dynamics or less reliable particle assignment due to signal-to-noise limitations. Classes having NU resolution worse than 4.5 Å are less reliably determined than the others. Specifics of the processing protocol (Li et al., 2022) are described in the text and the Methods.

**Figure. S5 (associated with Fig. 1):**
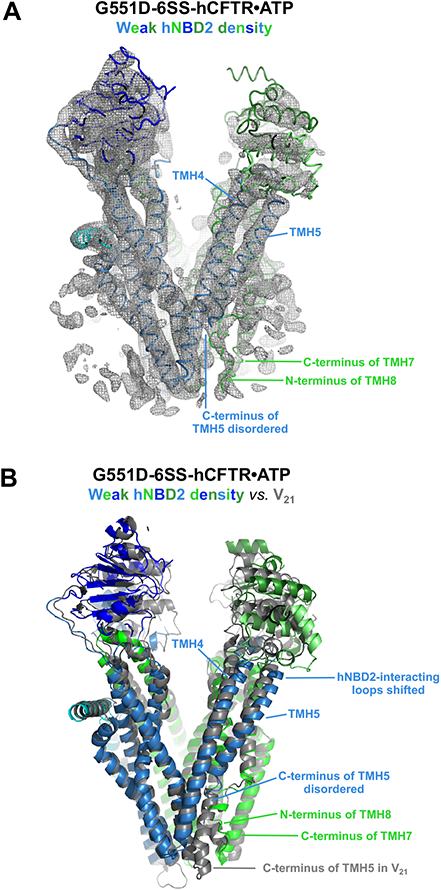
Structural differences in the weak hNBD2 density conformational class. **(A)** Cryo-EM density map for the weak hNBD2 density conformational class in the GCER for G551D-6SS-hCFTR/ATP/VX770 (Fig. 1). The density in the region of the extracellular exit pore is more consistent with the N-terminus of TMH8 occupying its standard position in the V_21_ and V_17_ structures and the C-terminus of TMH5 being disordered, but this density has features suggesting there could potentially be a mixture of that conformation and the conformation observed in the V_31_ structure (Fig. S6) in which the N-terminus of TMH8 is disordered and replaced at its standard position by the C-terminus of TMH5. **(B)** Ribbon diagrams showing the refined coordinate model for the weak hNBD2 density map in panel **A** superimposed on the V_21_ model based on least-squares alignment of the non-domain-swapped core of TMD2, with the weak hNBD2 density model colored according to domain organization as in Fig. 1A and the V_21_ structure colored gray.

**Figure. S6 (associated with Fig. 1):**
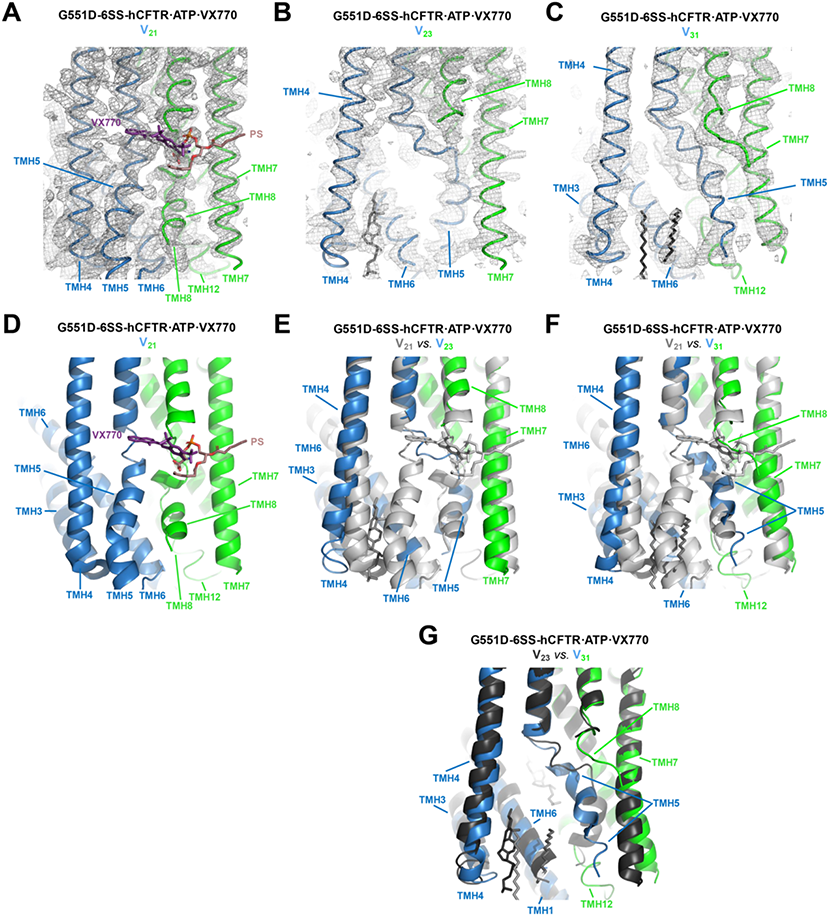
Conformational changes in the exit pore region of the TMDs in the conformational classes with wide TMD opening angles. **(A-C)** Cryo-EM density maps in the vicinity of the VX770 binding-site and extracellular exit pore for the V_21_, V_23_, and V_31_ conformational classes in the GCER for G551D-6SS-hCFTR/ATP/VX770 (Fig. 1). **(D-F)** Ribbon diagrams of the corresponding refined coordinate models (Table S1) colored according to domain organization as in Fig. 1A with the V_23_ and V_31_ models in panels **E**-**F** superimposed on the V_21_ model (gray) based on least-squares alignment of the non-domain-swapped core of TMD2. **(G)** Superposition of the V_23_ model (black) and V_31_ model (colored according to domain organization) based on least-squares alignment of the non-domain-swapped core of TMD2. In the V_23_ and V_31_ conformations, the N-terminus of TMH8 is largely disordered and displaced by the C-terminus of TMH5 from its standard position in the V_21_ and V_17_ structures.

**Figure. S7 (associated with Fig. 4):**
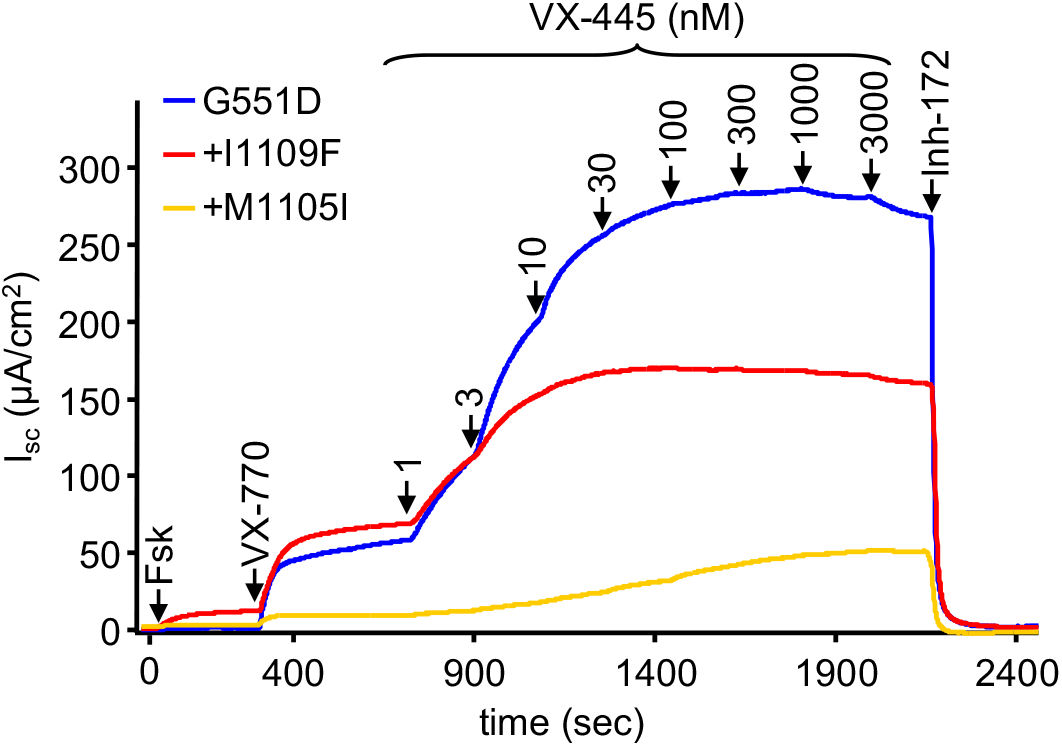
Short-circuit currents (Isc) in CFBE41o-cells expressing G551D-hCFTR variants harboring mutations in the VX445 binding site upon activation with forskolin and VX770 followed by potentiation with increasing concentrations of VX-445. These experiments were conducted in polarized monolayers mounted in an Ussing chamber using 20 µM forskolin (Fsk) and 3 µM VX770. The corresponding dose-response curves are shown in Fig. 4E.

